# Conservation units for anadromous Arctic Char (*Salvelinus alpinus*) in the Canadian Arctic informed by genetic structure, population connectivity and adaptive genomic variation

**DOI:** 10.1101/2025.10.18.683098

**Authors:** Xavier Dallaire, Anne Beemelmanns, Les Harris, Colin Gallagher, Ross Tallman, Jean-Sébastien Moore

## Abstract

Intraspecific genetic diversity is a crucial aspect of biodiversity conservation as it preserves evolutionary potential and enhances resilience to environmental change. Genomic-informed delineation of Conservation Units (CUs) offers ways of subdividing species into groups based on historical isolation and adaptive differentiation, to develop biologically relevant conservation and management policies. CUs have been defined in many species of harvested anadromous salmonids, but broad scale data remains lacking in the Canadian Arctic, where anadromous Arctic Char (*Salvelinus alpinus*) dominates catches in Indigenous-led subsistence and commercial fisheries. In this study, we use low-coverage whole-genome data from 30 Canadian populations of Arctic Char to define CUs based on population structure and connectivity, as well as adaptive genetic variation. We highlight two main genetic groups, each of which comprises three subgroups, or candidate CUs: the North (above the 67th parallel), including the North Baffin Island, Kitikmeot, and Inuvialuit Settlement Region CUs; and the South (below the 67th parallel), including the South Baffin Island, Ungava Bay, and Hudson Bay CUs. This delimitation is supported by areas of low effective migration between candidate CUs, as well as isolation-by-environment, which suggests adaptive differentiation. Finally, we discuss opportunities and caveats relating to linkage when identifying adaptive genetic variation from whole genome sequencing data through genome scans and Gene-Environment Associations.

## Introduction

Given the numerous threats to global biodiversity, there has been a call for conservation at the levels of ecosystems, species, and within-species genetic diversity (Laikre et al., 2020). Of these three levels, species conservation has gathered much of the attention, suggesting a simple categorization of nature and offering a common metric of biodiversity. In practice, the delimitation and definition of species is rarely straightforward (Mace, 2004; Frankham et al., 2012; Garnett & Christidis, 2017), and speciation proceeds via gradual genomic divergence among isolated populations driven by the combined effects of genetic drift and natural selection in contrasted habitats (Roux et al., 2016). Both these evolutionary forces are subject to fluctuations over long timescales, as populations reconnect in secondary contact zones, or changes in the environment shift selective pressures (Seehausen et al., 2008; Rosenblum et al., 2012). The resulting intraspecific genetic diversity is a key component of biodiversity that warrants protection, as it underpins evolutionary potential and enhances resilience to environmental change and anthropogenic pressures (Forest et al., 2007; Schindler et al., 2010).

Biological systems are inherently complex, and speciation is best understood as a continuous process. Nevertheless, discrete intraspecific classifications remain necessary to inform conservation priorities within limited budgets and to guide the management of harvested species. There exists a vast body of literature on the delineation of conservation units (CU), which generally relies on the identification of historical isolation (leading to genomic divergence) and adaptive differentiation (Waples, 1991; Moritz, 2002; Coates et al., 2018). Among the numerous CU models developed over the years, the Evolutionary Significant Unit (ESU; first proposed in Ryder, 1986) aims to provide a framework for the definition of subspecific groups that represent both unique genetic variation and the historical or ongoing connectivity among populations within a species. Depending on its definition, it can either be based exclusively on genetic characters or incorporate ecological data (Fraser & Bernatchez, 2001). As an additional concept, the Management Unit (MU) can be defined within ESUs as are demographically independent populations identified based on contemporary gene flow and population dynamics rather than deep evolutionary history. This subdivision allows managers to implement local actions, such as supplementation, restoration, or monitoring, that maintain genetic diversity, reduce inbreeding, buffer against environmental fluctuations, and support the long-term viability of the broader ESU (Moritz, 2002).

The concept of the ESU has been translated into legislation in multiple countries (e.g., the USA ‘Endangered Species Act’ and the Australian ‘Endangered Species Protection Act’). In Canada, the ‘Species at Risk Act’ applies a similar concept of Designatable Units (DU, Green, 2005), which are considered discrete if there is evidence of limited transmission of heritable traits or markers between them, or if they are separated by a natural disjunction in their distribution. A discrete unit is then considered significant if its evolutionary trajectory dates back long enough to create an evolutionary history not found elsewhere in Canada (e.g. units originating from distinct glacial refugia) or if it possesses unique and verifiable adaptive traits.

Genetic data has been central to work related to CUs, as their delineation often revolves around a two-step process: 1) describing the structure of populations, which is often hierarchical within species, and 2) choosing the level of distinctiveness appropriate for defining units, which implies some degree of subjectivity (Waples, 1995). Cytoplasmic (both mitochondrial and chloroplastic) DNA has long been used to characterize intraspecific lineages (Avise et al., 1987), while microsatellites have contributed to investigating population structure and connectivity (Ellegren, 2004). However, genomic data, i.e. genome-wide datasets produced by genotyping-by-sequencing (GBS, thousands of SNPs) or whole-genome sequencing (WGS, millions of SNPs), have the potential to improve delineation of CUs by quantifying adaptive variation in addition to neutral variation (Funk et al., 2012). This can be done by scanning the genome for loci displaying higher differentiation than expected under neutral evolution, suggesting the imprint of natural selection, or assessing correlations between allele frequencies and biologically relevant environmental variables (hereafter gene-environment associations, GEA, Rellstab et al., 2015; Dauphin et al., 2023).

Anadromous salmonids (i.e., those migrating from marine to freshwater habitats to spawn) represent a diverse group of fishes with significant economic and cultural importance worldwide. Their strong homing behavior (i.e., returning to natal sites for spawning, thus limiting gene flow among populations) plays a crucial role in shaping their population structure and is a factor long recognized as key for their conservation and management (Moulton 1939; Thompson 1959). Many salmonid populations are facing strong harvesting pressures and are actively managed within CUs to ensure sustainability and long-term population persistence. For instance, Atlantic Salmon (*Salmo salar*) along Canada’s eastern coast are currently managed within 15 extant DUs (COSEWIC, 2010; Moore et al., 2014). However, further subdivisions into 19 DUs was recently suggested based on genomic data (Lehnert et al., 2023). On the western coast, Pacific Salmons (*Oncorhynchus spp.*) are divided into dozens of CUs per species, with larger watersheds, such as the Fraser River in British Columbia, supporting multiple units (Xuereb et al., 2022; DFO, 2024). Overall, CUs have proven essential for guiding effective management and sustaining salmonid populations across diverse systems.

Northern Canada is home to populations of Arctic Char (*Salvelinus alpinus*), a salmonid whose anadromous form has long been the most harvested fish across Inuit Nunangat (i.e. the homeland of Inuit; Friesen, 2004; Priest & Usher, 2004). Arctic Char fisheries are predominantly small-scale and harvested for subsistence purposes (Roux et al., 2019; Tallman et al., 2019), but collectively amount to a high replacement value, i.e., the estimated cost of substituting the harvested fish with equivalent grocery-bought protein (Brubacher, 2004). While subsistence stocks are locally managed, some regions harbour commercial fisheries that are co-managed between the Indigenous land claim body and federal government (Fisheries and Oceans Canada). Continent-wide genetic variation in Arctic Char has been investigated using mtDNA and microsatellites. These studies suggest that most North American Arctic populations derive from a single High Arctic glacial refugium, the Arctic lineage (Moore et al., 2015), with the exception of Labrador (Salisbury et al., 2019) and Greenland (Jacobsen et al., 2022), where an additional Atlantic lineage is also present. However, recent whole-genome data revealed that admixture between the Arctic and Atlantic lineage was a prominent feature of Arctic Char genomes in all Canadian populations south of the 67^th^ parallel, redefining the extent of the second contact between lineages (Dallaire et al., 2025). Other recent genomic datasets for the species were focused on spatial scales from local to regional and have been used to investigate questions relating to population structure and connectivity (Moore et al., 2017; Li et al., 2021), morphotype differentiation (Kess et al., 2021), and local adaptation (Madsen et al., 2019; Dallaire et al., 2021; Layton et al., 2021).

In this study, we revisit the whole-genome resequencing dataset introduced in Dallaire et al. (2025), including more than 1,000 anadromous Arctic Char whole-genomes in 30 sampling sites across the Canadian Arctic, to define candidate CUs at a national scale. To inform these units, we 1) characterize in depth the hierarchical genetic structure of populations in relation to administrative regions in the Inuvialuit Settlement Region (ISR, Northwest Territories), Nunavut, and Nunavik (Québec), and 2) identify areas of lower gene flow. Then, we aim to 3) disentangle patterns of i) isolation-by-distance, ii) isolation-by-environment, and iii) isolation-by-colonization (related to post-glacial recolonization) that characterize this structure. Finally, we 4) detect signals of natural selection and explore patterns of population structure driven by putatively adaptive genetic variation using genome scans and Gene-Environment Associations. We discuss those results through the lens of the Designatable Unit framework and explore and discuss the added value of whole-genome data to conservation genomics.

## Methodology

### Sequencing and SNP calling

We reanalysed sequencing data from 1016 Arctic Char from 30 Canadian populations, published in Dallaire et al. (2025). Briefly, whole-genome sequences were obtained following Therkildsen & Palumbi (2017) on DNA extracted from fin clips from fish harvested at the mouth of rivers in the Inuvialuit Settlement Region (Northwest Territories), Nunavut, and Nunavik (Québec) between 2007 and 2023 (Fig.1a). As in Dallaire et al. (2025), data were processed following the pipeline described at https://github.com/enormandeau/wgs_sample_preparation: sequences were trimmed and aligned on the ASM291031v2 reference genome (likely from *Salvelinus malma* or a *S. alpinus* x *S. malma* hybrid; Christensen et al., 2021), duplicate reads were removed, indels were realigned and overlapping ends of paired reads were clipped.

Mapped and cleaned reads yielded an individual depth of coverage around 2X, and we estimated Genotype Likelihoods (GL) at polymorphic positions identified in Dallaire et al. (2025). Those positions were distributed on all assembled linkage groups, but SNPs in sex-linked regions identified in Beemelmanns et al. (2024) were removed from all analyses. The SNP list was filtered for paralogs and other deviations from expected patterns due to transposable elements and ancestral autoploidization in salmonids, using *ngsparalog* (Linderoth, 2018) following Dallaire et al. (2023). We kept SNPs with global minor allele frequency above 1% and with a minimal depth of coverage of 1X in at least 75% of individuals. We used *ngsLD* (Fox et al., 2019) to estimate linkage between SNPs within 500 kb using a random subset of half the samples to create a LD-pruned dataset by iteratively removing SNPs with the graph method until no SNP pairs within 200kb had an r^2^ above 0.1.

### Extraction of environmental data

We used the ArcGIS software v10.4 (ESRI, 2011) to extract environmental data. Marine variables represent sea-surface values for factors of potential biological importance for Arctic Char available in the BIO-Oracle v3.0 dataset (Assis et al., 2024) and were averaged in a 20 km radius around each sampled river mouth and within 5 km from the coast. This standardized approach was used to characterize the local coastal environment in accordance with current knowledge of Arctic Char marine habitat use in Arctic Canada (Spares et al., 2015; Moore et al., 2016; Harris et al., 2020). We selected annual coastal averages for five marine variables (temperature, salinity, primary productivity, dissolved O_2_, and ice cover) as they represented good predictors for values in summer, when Arctic Char occupies coastal habitats.

Freshwater variables were extracted from the RiverATLAS dataset by selecting the linear segment closest to the mouth of each sampled river, as these segments include values averaged over the entire upstream watershed. We initially considered seven variables: % of the watershed covered by lake or tree, average slope, sand fraction in the soil, and annual averages of discharge, air temperature, and precipitation. These variables are linked to both migratory challenges (e.g., slope, discharge, temperature) and spawning habitats (e.g., lake cover, sand in soil), two factors that may play an important role in local adaptation of Arctic Char (Moore et al., 2017; Dubos et al., 2023). Precipitation was later excluded because it was highly correlated with air temperature (Pearson r = 0.84), resulting in a final set of 11 environmental predictors (Table S1, Table S2, Fig. S1).

### Population structure

When describing population structure in the dataset, we used a priori administrative regions to categorize sampling sites across the study area, namely the Inuvialuit Settlement Region (Northwest Territories); Kitikmeot, Kivalliq and Baffin (Nunavut); and Nunavik (Québec). We used the term Baffin instead of Qikiqtaaluk, as this administrative region also includes the High Arctic Archipelago, which was not sampled as part of this study. Note that anadromous populations of Arctic Char are less common at those latitudes where resident and landlocked Arctic Char dominate (Reist et al., 2013). North and South Baffin were further split around the 67^th^ parallel. Note that the Pamiurluk Lake sampling site (PAM) was grouped within the Kivalliq region, despite technically being in Qikiqtaaluk, as this fish population is traditionally harvested by the people of Naujaat, a community in the Kivalliq region.

The overall distribution of genetic variation and population structure was initially evaluated through a principal component analysis (PCA) conducted on the LD-pruned dataset using PCAngsd 1.10 (Meisner & Albrechtsen, 2018). Subsequently, we examined the hierarchical population structure by applying NGSadmix (Skotte et al., 2013), estimating individual admixture proportions while varying the number of ancestral populations (*K*) from 1 to 30 (i.e. the number of sampling sites). For each value of *K*, we repeated the analysis over 50 independent runs, then used the CLUMPAK algorithm (Kopelman et al., 2015) to cluster runs with a similarity score over a fixed threshold of 0.85. We interpreted the cluster with the most runs (hereafter the major mode) for each value of K. Ancestry proportions were averaged over runs in the major mode and visualized using a stacked bar plot. To compare the fit of the model at different values of K, we 1) compared the proportion of runs in the major mode and 2) computed the Δ*K* (Evanno et al., 2005) by dividing the absolute value of the second order rate of change in the mean likelihood by the standard deviation of likelihoods for all runs of a given *K*.

Next, we used ngsDist (Vieira et al., 2016) on the unpruned dataset to compute genetic distances between every individual pair. We chose the uncorrected p-distance, i.e. the proportion of nucleotide sites (polymorphic in the dataset) at which they differ. SNPs with missing data in either individual of the pair were ignored. To express p-distances as the proportion of genome-wide differences between two individuals, we multiplied them by the ratio of the number of SNPs (4,044,588) to the total number of sites in the reference genome (1,516,033,356). We then built an unrooted neighbour-joining tree on the distance matrix using the BIONJ algorithm (Gascuel, 1997) in the ape v5.7 R package (Paradis & Schliep, 2019). We visualized the tree with the R packages *ggtree* (Yu et al., 2017) and *ggtreeExtra* (Xu et al., 2021), while collapsing monophyletic groups of individuals from a single sampling site, then aligned individual ancestry proportions estimated with ngsAdmix for K = 2, 7, and 30 genetic clusters with the corresponding tree tips. These values of K were chosen as they represented Arctic vs Atlantic ancestry as detailed in Dallaire et al. (2025) (K = 2), roughly corresponded to *a priori* geographic regions (K = 7), or matched the number of sampling sites (K =30).

### Connectivity, Isolation-by-distance, Isolation-by-environment

Dallaire et al. (2025) revealed strong patterns of isolation-by-distance that were distinct in groups of populations above (hereafter northern) and below (southern) the 67^th^ parallel, with southern population pairs being more genetically distant at equal marine distances. Here, we revisited the relationship between genetic distance, as measured by the pairwise fixation index (F_ST_) and allele frequency difference (AFD), and either marine, environmental, and ancestry distance to investigate the interplay between i) isolation-by-distance (IBD), ii) isolation-by-environment (IBE), and iii) isolation-by-colonisation (IBC; defined as the proportion of genetic distance explained by the difference in genetic background contributions from two glacial lineages that recolonized the Canadian Arctic), respectively.

F_ST_, AFD, and marine distances were estimated as in Dallaire et al. (2025). Environmental distance between sites was calculated by performing a PCA on all environmental variables and measuring the Euclidian distance between population coordinates on the first seven axes, which encompassed 92.2% of the variation. Ancestry distance was measured as the difference in the average ancestry proportion for a sampling site in the NGSadmix analysis for K = 2, to represent a proxy of the proportion of ancestry from the Arctic and Atlantic lineages.

The correlations between each measure of genetic distance and either marine, environmental, or ancestry distances was evaluated separately using simple Mantel tests as implemented in the R package *vegan* (Oksanen et al., 2022). We then used partial Mantel tests to test whether each correlation was robust to controlling for either other type of distance (e.g. AFD ∼ marine distance while controlling for ancestry distance, then for environmental distance, etc.). To control for the effect of two distance matrices conjointly, we first built a linear model between the genetic distance and the admixture model while accounting for the category of the population pair (i.e. two northern populations, two southern populations, or one northern and one southern population), and used the residuals as a substitute for genetic distance in a partial Mantel test with marine or environmental distance. To further investigate which component of the environment could contribute to isolation-by-environment, we repeated partial Mantel tests using the difference in the value for each environmental variable as environmental distance. We compared Mantel r statistics and their associated p-values estimated from 999 permutations for tests including all populations pairs, then only northern or southern pairs. We excluded GEO from all Mantel tests due to its high genetic distance from other sites. These diverged from all patterns of isolation, likely due to heavy introgression from the Atlantic lineage in genomes from this population (Dallaire et al., 2025).

To characterize geographic areas of limited connectivity, we conducted an Estimated Effective Migration Surfaces (EEMS) analysis (Petkova et al., 2016) using FEEMS (Marcus et al., 2021). Briefly, the effective migration rate is comparable to the concept of effective population size (N_e_), in that it represents the rate of migration at which an idealized stepping stone model at equilibrium would produce genetic dissimilarities equivalent to values observed in the data. In FEEMS, we modified the python scripts *spatial_graph* and *run_cv* to skip the allele frequency estimation from genotypes and used the population MAF estimated in ANGSD as input since we did not call individual genotypes. We used the R package *dggridR* (Barnes & Sahr, 2024) to create a triangular grid at resolution 7 (∼ 55km between cells) over the study area. To select the value for the smoothing parameter λ, we followed the leave-one-out cross-validation method suggested in Marcus et al. (2021): we repeated the EEMS analysis after removing one population at a time with 20 values of λ varying from 0.1 to 10,000 and compared the migration surfaces for λ displaying local minima in cross-validation errors.

### Loci under putative selection and gene-environment association

SNPs with a global minor allele frequency over 0.05 were investigated for signatures of selection using *pcadapt* (Luu et al., 2017) and the XtX statistic in *Baypass v2.1* (Gautier, 2015), and for Gene-Environment Associations (GEA) using the Bayes Factor statistic in *Baypass* and redundancy analyses (RDA).

We applied the *pcadapt* procedure in PCAngsd, which allows for individual genotype likelihoods as input. Briefly, *pcadapt* scans the genome using significant components of a PCA, here the first 15 PCs, as selected by PCAngsd. The multi-dimensional z-scores in output were converted to Mahalanobis distances using the orthogonalized Gnanadesikan–Kettenring method, corrected for genomic inflation following recommendations (Luu et al., 2017), then converted into p-values via a χ^2^ test with 14 degrees of freedom. To build the input for *Baypass*, we converted population allele frequencies estimated in ANGSD (-domaf 1) to allele counts by multiplying (then rounding) the frequencies by the SNP-specific number of individuals to account for missing data. We first sampled 10,000 random SNPs to compute a covariance matrix with the core model in *Baypass*, then used this matrix in the standard covariate model with every SNP (maf > 0.05) and all 11 environmental variables. We ran five replicate analyses and used the median XtX as an index of selection and the median Bayes Factor (BF) as a measure of the association between the allele frequency of a SNP and each environmental variable.

As a second method to detect Gene-Environment Associations, we used redundancy analyses, which are constrained ordinations that assess the relationship between a multivariate response variable (here allele frequencies in sampling sites) and explanatory variables (the environmental factors). To account for collinearity between variables, we ran a first RDA with all variables in the R package *vegan*, then iteratively removed the variable with the highest Variance Inflation Factor (VIF) until all VIF were under 10 (we only removed Dissolved O_2_ this way). We then ran a backward selection of model with the function *ordistep*, and iteratively pruned variables from the model until all variables contribute to the model according to an ANOVA-like permutation test (p < 0.1). Finally, we assessed the significance of RDA axes through an ANOVA-like permutation test on the selected model and kept axes with p < 0.05.

Since the ancestry proportion from the Arctic and Atlantic lineages (as estimated by the NGSadmix analysis for K = 2; see also Dallaire et al. 2025) was a major determinant of the genetic variation in our study area and strongly correlated with some of the environmental variables, we also conducted a partial RDA (pRDA) using the average ancestry proportion as a covariate. We followed the same variable and axes selection steps as described above. We use the *varpart* function in the R package *vegan* to estimate the portions of the genetic variation that was explained conjunctly and distinctively by environment and lineage ancestry in the final pRDA.

Every selection scan and GEA methods described above produced summary statistics by SNP that were aggregated by gene using a windowed Z analysis (WZA; Booker et al., 2024). WZA combines the information of multiple SNPs linked in a genomic region to assess if this region is in GEA or displays signs of selection. Since linkage varies across the genome, and in the absence of a robust recombination map, we opted to use genes annotated on the reference genome instead of windows of an arbitrarily defined length. We ran separate WZAs by using as summary statistics i) the *p*-values from pcadapt, ii) the XtX and iii) BF (for each environmental variable) from Baypass, and iv) the squared loading on each significant axis of the RDA and pRDA. Briefly, those statistics were ranked, ordered and converted to empirical p-values, then to z-scores that were combined for SNPs inside each gene (i.e. between the start and end position based on the annotated reference genome) to compute a *p*-value corrected for the number of SNPs in the gene (Booker et al., 2024).

For each WZA, we listed top candidate genes (i.e. the genes with p-values < 0.001), thus showing the strongest signal of selection or association with the environment. To explore the relationship between our top candidate genes and regions of low recombination as inferred by local PCA in Dallaire et al. (2025), we conducted a series of χ^2^ tests to check if top candidates identified by each analysis were more likely than by chance to be found in long putative local ancestry tracts, then in other local PCA outlier regions not correlated with the admixture gradient between the Arctic and Atlantic (as listed in Table S3).

For each selection scan or GEA analysis, we performed a gene ontology (GO) enrichment test with goatools 1.2.3 (Klopfenstein et al., 2018) on all top candidate genes, and then on genes identified as top candidates by more than one method. We used a custom annotation table, obtained by mapping Arctic Char transcripts (accession GCF_002910315.2; ASM291031v2) against the Swissprot 1.2 database (UniProt Consortium, 2024) with blast 2.12.0 (Camacho et al., 2009), according to the pipeline GAWN v0.3.5 (https://github.com/enormandeau/gawn). As we lacked power to detect selection in WZA for genes with few SNPs, only genes on which we considered at least 5 SNPs were kept as the background set. Terms for biological processes (BP) with corrected p-values (Benjamini & Hochberg, 1995) under 0.05 were considered significantly enriched.

## Results

### Genetic population structure

Out of the 5.98 million polymorphic SNPs in Canadian and Greenlandic Arctic Char populations identified in Dallaire et al. (2025), 4.04 million passed the coverage filters and displayed MAF above 0.01 in Canadian populations alone. Subsequent LD-pruning reduced this number to 277,570 SNPs. To facilitate knowledge transfer to local policy makers, populations were first categorized *a priori* according to administrative regions of Inuit Nunangat: the Inuvialuit Settlement Region (ISR, Northwest Territories); Nunavik (Québec); and Kitikmeot, Kivalliq, and Baffin (Nunavut) (Fig. 1a, circles and labels).

**Figure 1:**
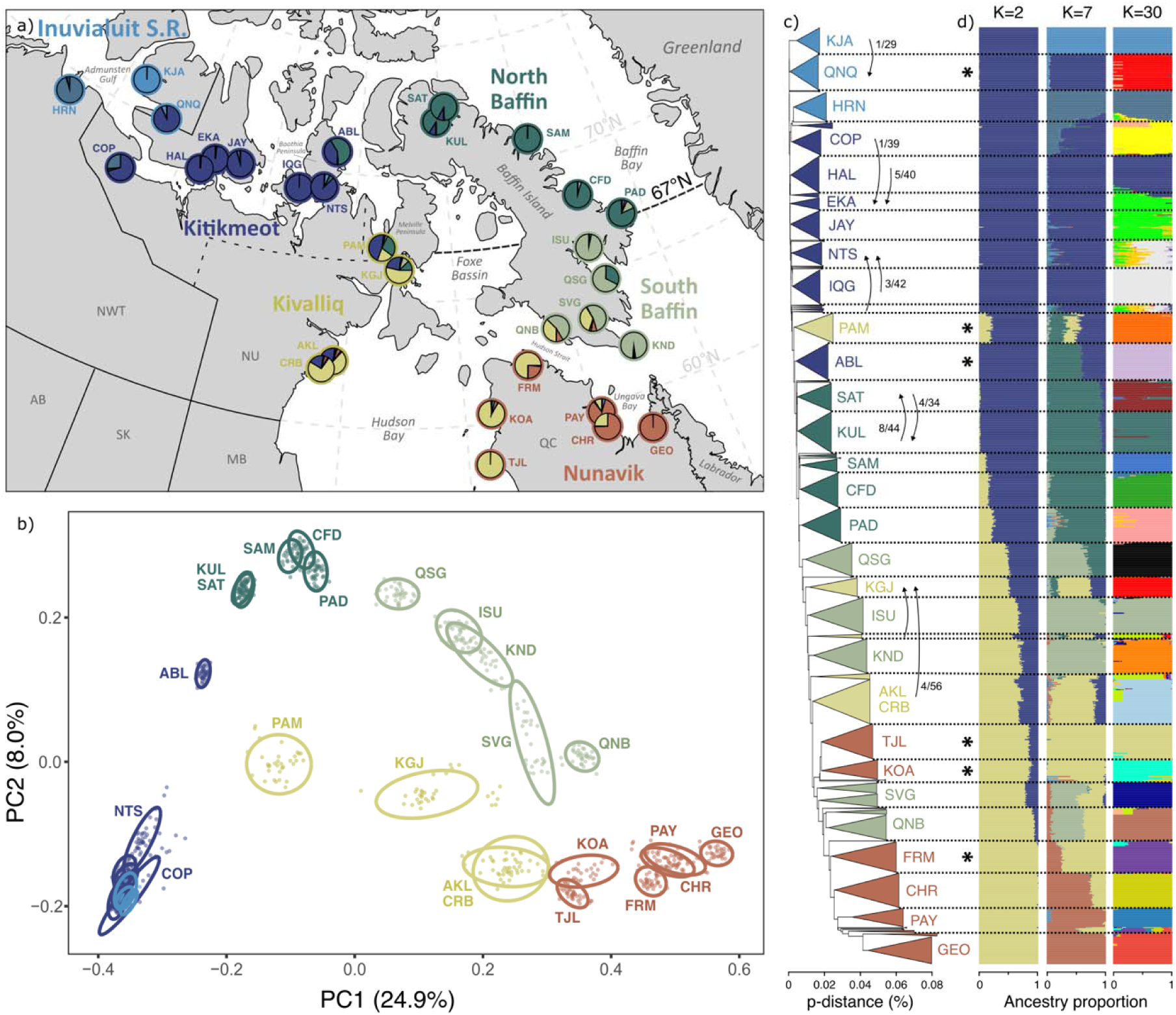
a) Arctic Char sampling sites with colored labels showing their administrative region of origin. Pie charts show the site average in ancestry proportion for K = 7 genetic clusters, as detailed in panel d). b) The first two axes of a principal component analysis for 277,570 independent SNPs. 95% confidence interval ellipses are drawn around each site and the percentage of variance explained by each axis is noted in parentheses. c) Unrooted neighbor-joining tree built on genome-wide uncorrected p-distances between Arctic Char samples. Monophyletic clades where at least 80% of samples originate from a single site (or a pair of neighbouring sites) are collapsed as triangles and truncated at the minimal length. Sampling sites are delimited by dashed lines and arrows note the origin of samples in smaller clades. Arrows accompanied by fractions denote clades where a portion of individuals were captured at a different site. d) Individual proportion of ancestry for K = 2, 7, and 30 genetics clusters, CLUMPAK-averaged over replicate runs in the major mode (out of 50 runs). Individuals are aligned to their position in the tree in c). Note that this figure reuses the same color palette for two codes: administrative regions in panel a) (labels, circles), b), and c); and their most concordant genetic cluster (K = 7) in panel a) (pie charts) and d). Discordance between administration regions and genetic clusters are noted by an asterisk. Additional colors for the K = 30 plot were assigned randomly.

A principal component analysis revealed a population structure consistent with geography on the first two axes (comprising 32.9% of the variation; Fig. 1b). However, most sites in ISR and Kitikmeot were densely clustered on these axes and while HRN and KJA were distinct on the fourth axis (3.6%), the rest of north-western sites remained indistinguishable near the middle of the plot for the first 10 axes (totaling 55.8% of the variation; Fig. S2). A neighbour-joining tree based on p-distances displayed a similar geographic pattern, with north-western individuals on shorter branches than south-eastern individuals (Fig. 1c). Many sampling sites clustered into exclusive groups, with the notable exception of closely located pairs of sites: EKA and HAL (Kitikmeot; 39 km, F_ST_ = 0.010, Table S4), AKL and CRB (Kivalliq; 64 km, F_ST_ = 0.011), and SAT and KUL (North Baffin; 65 km, F_ST_ = 0.014). Interestingly, PAY and CHR were only marginally further apart (69 km), but exhibited much stronger structure (F_ST_ = 0.078). Finally, some sampling sites displayed considerable substructure, as their clade was split into subgroups (e.g., SVG, KUL-SAT) or had a few individuals grouping into smaller distant clades (e.g., NTS, KGJ).

The *NGSadmix* analysis of ancestry proportions with *K* ranging from 2 to 30 ancestral populations revealed a highly hierarchical structure (Fig.1d, Fig. S3, Fig. S4). At K = 2, eastern Nunavik and northern sites displayed pure ancestry for different groups, and sites in between followed a cline in admixture closely matching the pattern attributed to Arctic vs. Atlantic lineage ancestry in Dallaire et al. (2025). At K = 30, the analysis discriminated most sampling sites, in line with clades identified in the neighbour-joining tree. Interestingly, K = 7 groups produced a structure parallel to geographic regions defined *a priori* (Fig. 1a, pie charts), with a few key discordances. For example, PAM clustered more closely with Kitikmeot than Kivalliq populations, and ABL aligned with North Baffin rather than Kitikmeot. Most notably, at this level of structure, populations along Hudson Bay and Ungava Bay clustered into different genetic groups, rather than aligning with the Kivalliq and Nunavik regions defined *a priori*. We hereafter refer to groups of sampling sites sharing a dominant genetic cluster at K = 7 as our proposed CUs, with the exception of HRN and KJA who were grouped into the Inuvialuit Settlement Region CUs, as discussed later.

### Connectivity and isolation

Pairwise F_ST_ ranged from 0.011 (HAL-EKA) to 0.407 (IQG-GEO), while average allele frequency difference (AFD) in SNPs with MAF > 0.01 ranged from 0.033 (HAL-EKA) to 0.231 (HRN-GEO) (Table S4). The strongest predictor of genetic distance was lineage ancestry difference, with Mantel correlations (r) of 0.92 for F_ST_ and 0.90 for AFD (Fig.2a, Table S5). In comparison, marine geographical distance showed lower correlations (r = 0.65 for F_ST_, 0.58 for AFD), and environmental distance (summarized by PCA of 13 factors) had even weaker, but still significant (p < 0.01), correlations (r = 0.49 for F_ST_, 0.51 for AFD) (Table S5. At equal ancestry, marine, or environmental distances, pairs of populations from the southern group were more genetically distant than northern pairs, and the George River (GEO) was strikingly more distant from all other populations than other southern sites.

Partial Mantel tests indicated no signs of isolation-by-distance in the global dataset when correcting for both ancestry and environmental distance (Fig.2b, r = −0.02, p = 0.972), but signs of weak isolation-by-environment when correcting for ancestry and marine distance (Fig. 2c, r = 0.23, p = 0.025). When repeating these analyses exclusively for northern or southern pairs, we revealed apparent isolation-by-distance in the North (Fig. 2b, r = 0.52, p < 0.001), and isolation-by-environment in the South (Fig. 2c, r = 0.57, p < 0.001). Partial Mantel tests with specific environmental variables indicated that difference in discharge (r = 0.33) and primary productivity (r = 0.46) explained a significant portion (p < 0.01) of the variation in study-wide AFD, while differences in primary productivity (r = 0.47) and salinity (r = 0.42) explained a significant portion of the variation in AFD in southern pairs of sites (Table S5, Fig. S5).

**Figure 2:**
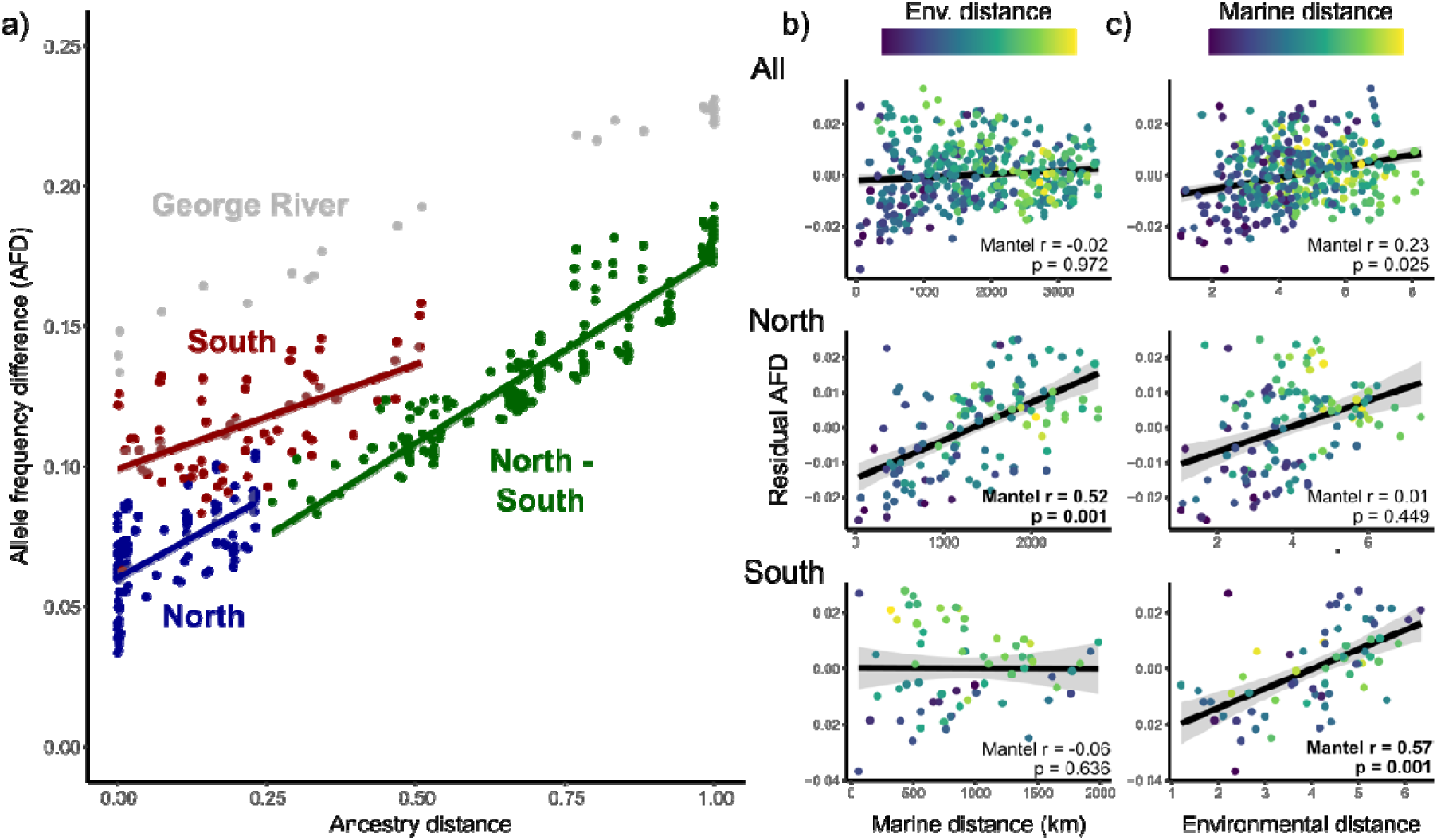
a) Linear regression of allele frequency differences (AFD) between a pair of populations and their ancestry distance (i.e. the difference in their average ancestry proportion, NGSadmix, K=2). Pairs were categorized depending on if they include two northern (blue), two southern (red), or one northern and one southern population (green). Pairs including the George River (GEO) are displayed in grey and excluded from panel b-c. Partial Mantel tests were conducted between the residuals of the model from panel a and either b) marine or c) environmental distance for all populations (top), or either northern (middle) or southern ones (below). The remaining distance, i.e. b) environmental or c) marine, was used as covariable and is displayed in color (yellow to blue). Mantel r statistics are noted in bold when the associated p-value was under 0.01.

We estimated effective migration rates across the study area from allele frequencies. The resulting migration map at different values of the smoothing parameter lambda mainly highlights an area of low effective migration between the previously described northern and southern regions (Fig. 3, Fig. S6). Smaller areas of low effective migration are observed over the Boothia Peninsula (between NTS and ABL), Amundsen Gulf (between HRN, KJA), Hudson Strait (between QNB and FRM), and inland Nunavik (between Hudson Bay and Ungava Bay). Sampling sites within CUs were generally connected by areas of effective migration rates higher than 1.

**Figure 3:**
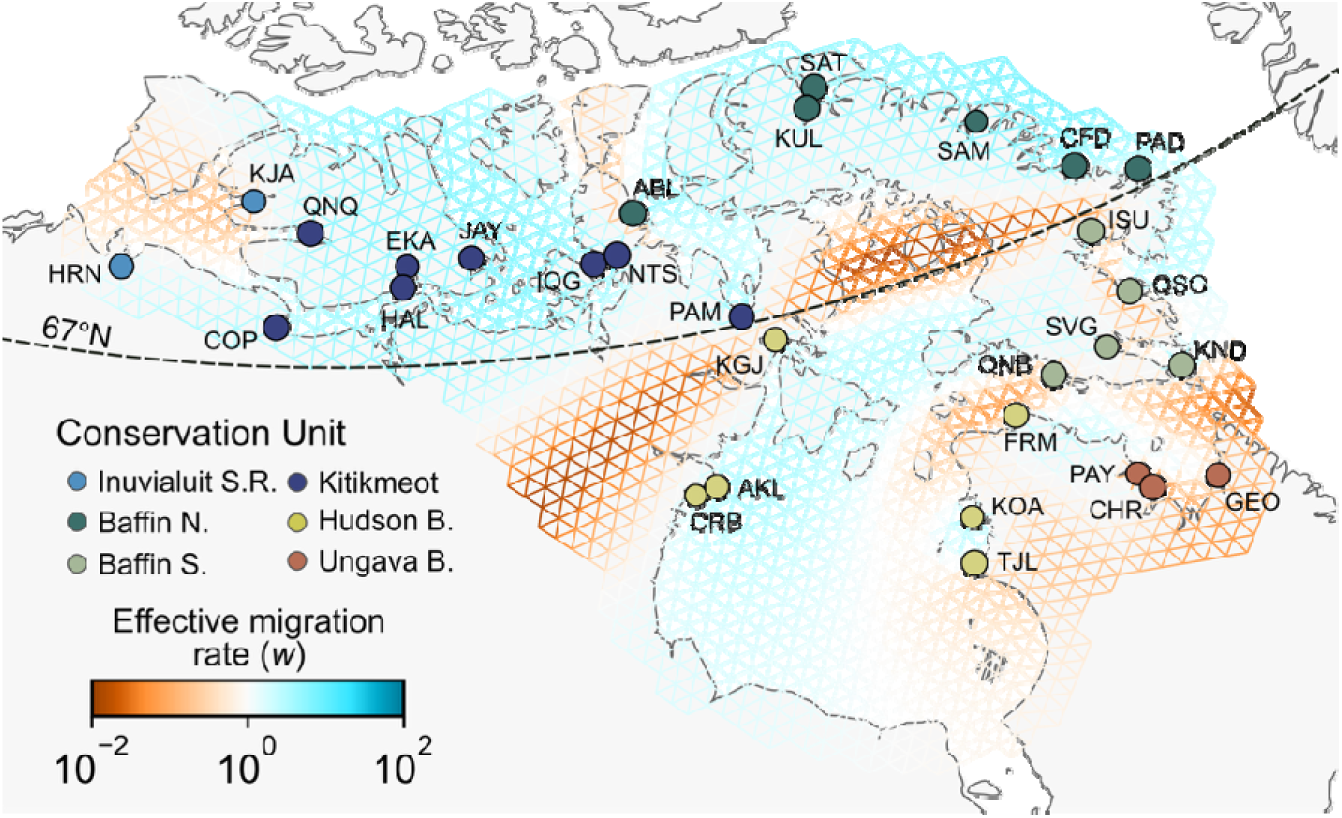
Edge-specific effective migration rates (w) estimated from sampling allele frequencies in FEEMS with a smoothing parameter λ = 21.5. Sampling sites were repositioned to the nearest node on a triangular grid (cell spacing ≈ 55 km) and colored with the proposed conservation unit, following the most dominant genetic cluster in an NGSadmix analysis (K = 7), except for Inuvialuit S. R. (see Discussion for details).

### Selection scan and Gene-environment association

We investigated 2.6 million SNPs with MAF over 0.05 for signatures of selection, via pcadapt and the XtX statistic in Baypass (details below); or their association with environmental variables, with redundancy analyses without (RDA) and with (pRDA) correction for lineage ancestry, and the Bayes Factor statistic in Baypass.

A global RDA with all 11 selected environmental factors and Arctic-Atlantic lineage ancestry (as estimated by the average ancestry proportion inferred with NGSadmix at K=2) explained 62.4% of the variation in MAF between sampling sites (Fig. 4a). Environmental factors explained 53.3% of the variation, but a large part (38.4%) was conjointly explained by lineage ancestry, as further explored by the following RDA and pRDA.

**Figure 4:**
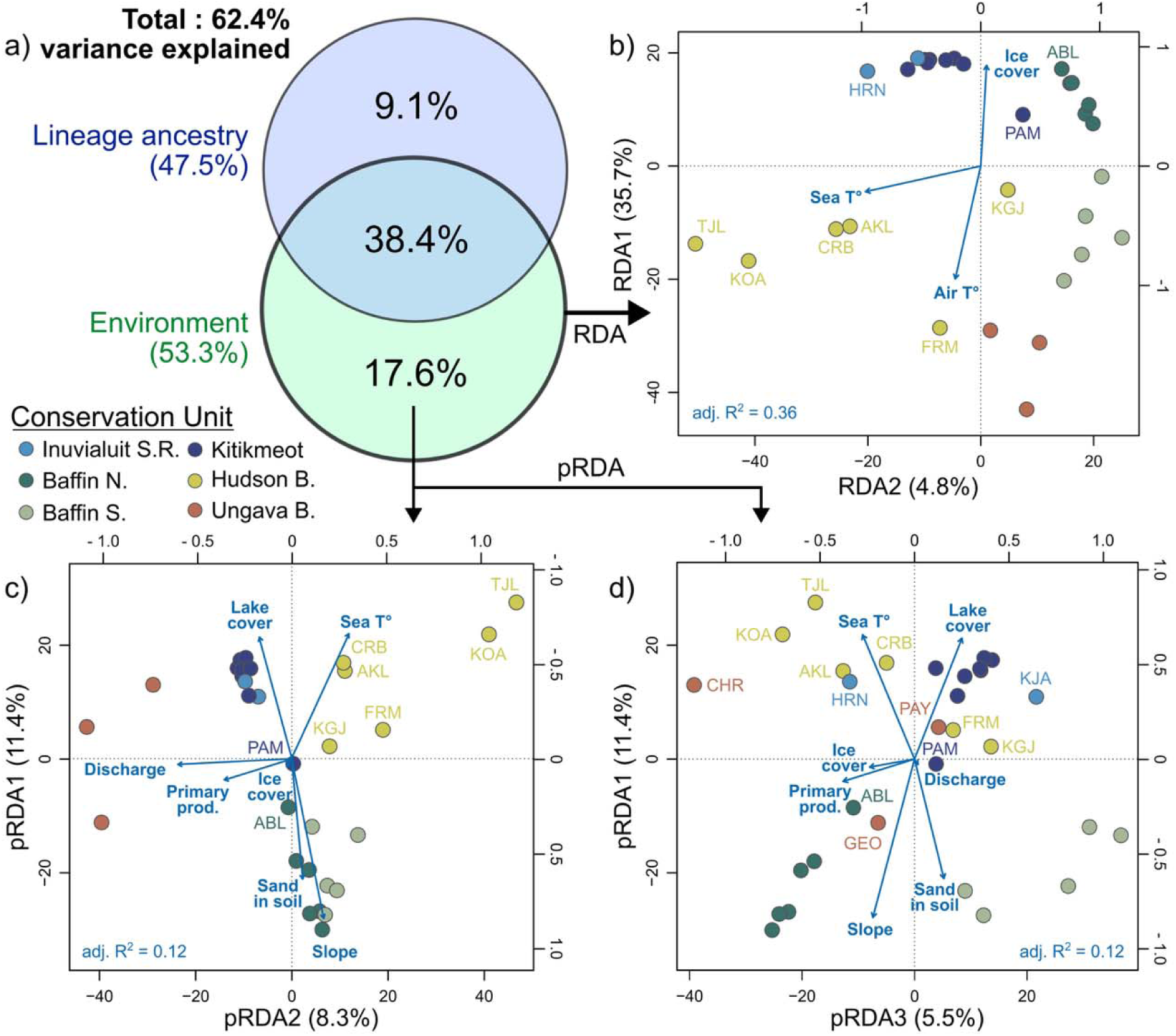
a) Proportion of the variation in allele frequency explained independently and conjointly by a set of 11 environmental variables and glacial lineage ancestry, as estimated by NGSadmix (K = 2). Biplots of b) a redundancy analysis (RDA, 2 significant axes) and c-d) a partial RDA accounting for the variation in ancestry (pRDA, 3 significant axes). Sampling sites are represented by points color-coded by their proposed conservation unit, and vectors represent contributing environmental predictors (retained after model selection) according to the scales on top and right axes.

After backward variable selection with ordistep, the first two axes of the RDA (adjusted R^2^ = 0.36) were significant (p < 0.05) and were largely correlated with 1) air temperature and ice cover; and 2) sea surface temperature (Fig. 4b). The first three axes of the pRDA (adjusted R^2^ = 0.12) were significant and were correlated with 1) sea-surface temperature, lake cover, fraction of sand in soil, and slope; 2) discharge and primary productivity; and 3) primary productivity and ice cover (Fig 4c-d). RDA1 was highly correlated with latitude and lineage ancestry, while RDA2 and pRDA1 discriminated Baffin Island sites from the rest of the study area. Both RDA2 and pRDA2 also highlighted the genetic and environmental distinction between western (Hudson Bay: TJL, KOA) and eastern (Ungava Bay: PAY, CHR, GEO) Nunavik.

With a windowed Z analysis (WZA), we combined SNP-specific statistics from the 5 methods (a total of 18 tests: pcadapt, XtX, 2 RDA axes, 3 pRDA axes, and 11 univariate tests for the BF statistics in Baypass) for 21,379 genes (with at least 5 SNPs) to provide rank-ordered support for selection or GEA, in the form of empirical p-values (Fig. 5a). The genome scan methods (pcadapt and XtX) identified respectively 151 and 188 top candidate genes (p < 0.001), with significant overlap (81, or 31.2% of genes in common, Fig. 5b). The GEA methods (BF, RDA, and pRDA) identified 2,079, 381, and 544 top genes. BF had notable overlap with the other methods: 164 (7.1%) of top genes were in common with the RDA and 250 (10.5%) with the pRDA (Fig. 5b). The RDA and pRDA had a similar level of overlap, with 57 top genes in common (6.6%). Top candidate genes from the XtX, pcadapt method or associated to pRDA1 or slope had more chance to be found in putative local ancestry tracts identified in Dallaire et al. (2025), while other local PCA outlier regions were enriched in pcadapt, RDA2, pRDA1, Lake cover, and Sea T° top candidate genes (χ^2^ test, adjusted p < 0.05; Table S6).

**Figure 5:**
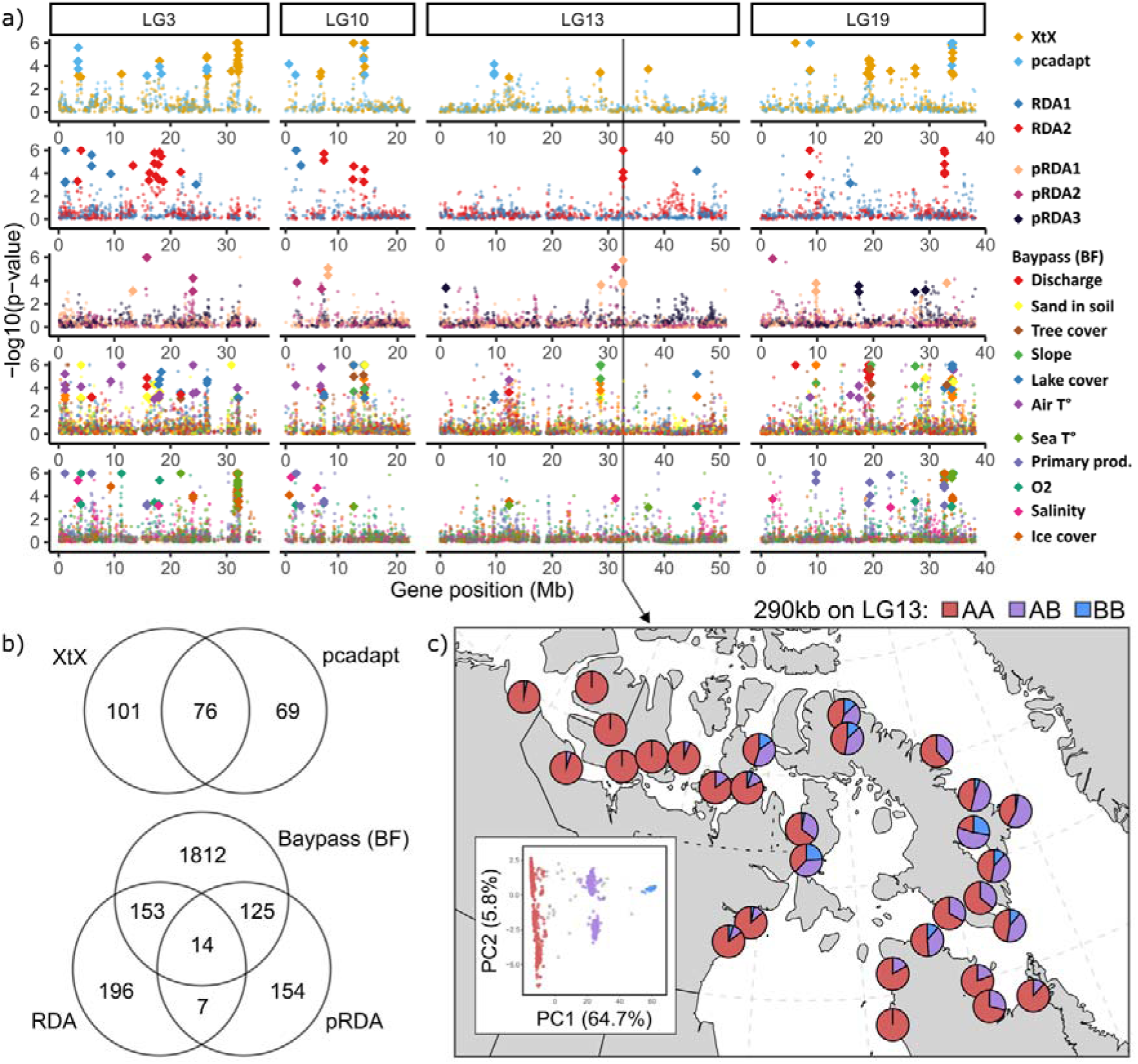
a) Support for selection scans or gene-environment associations on genes with at least 10 SNPs on 4 linkage groups, in 5 panels regrouping methods: 1) XtX (Baypass) or pcadapt; 2) redundancy analysis axes (RDA); 3) partial RDA axes; and Bayes Factors (BF, Baypass) for 4) freshwater and 5) marine variables. Support at the gene level (empirical p-values) was derived from combined per-SNP summary statistics in a windowed Z analysis (WZA). Top candidate genes (p < 0.001) for more than one type of test were highlighted with bigger diamonds than the other genes (small dots). b) Venn diagrams presenting the number of top candidate genes (p < 0.001) for selection (top) or gene environment association (bottom) and the overlap in genes between methods. c) Karyotype frequencies per sampling site for a 290kb-long putative inversion on LG13, as inferred by local PCA in Dallaire et al. (2025). The genomic position of the putative inversion is noted by an arrow in panel a.

The higher number of top genes identified by the BF method is linked to the univariate nature of the method, as top-ranking candidate genes were separately identified based on their correlation with each of the 11 environmental variables. Individual variables had on average 262.1 (sd = 14.7) associated top genes (empirical p < 0.001 in the WZA). Out of 2,079 unique BF top genes, 571 (27.4%) were associated with more than one variable (404 with two variables, 120 with three, 47 with four or more). Salinity and dissolved O_2_ (55 top genes in common), followed by lake cover and slope (19 genes), were the most frequent pairs.

Across all genome scans and GEA methods, 549 out of 2,634 unique top candidate genes (20.8 %) were identified by more than one method (Table S7, Fig. S7). We tested for enrichment in gene ontology (GO) terms in lists of outlier genes from each individual test, as well as the list of genes that were outliers according to at least two methods and identified 89 unique GO terms with a p-value for enrichment under 0.001 and associated with at least three top candidate genes (Table S8). However, after correction for multiple testing, no GO terms presented corrected p-values under 0.1.

## Discussion

In this study, we aimed to define conservation units for anadromous populations of Arctic Char across its continuous distribution in the Inuvialuit Settlement Region, Nunavut, and Nunavik in Arctic Canada. Through analysis of population structure and connectivity, we highlight two major population groupings, respectively above and below the 67^th^ parallel, which we interpret as having distinct evolutionary trajectories and reduced gene flow. These northern and southern groups were further subdivided in six candidate CUs (see Fig. 3) based on genetic similarity and connectivity. While we observed interconnected patterns of isolation-by-distance and isolation-by-environment across the landscape, glacial lineage ancestry was the strongest predictor of genetic distance among populations. Using genome scans and GEA, we detected a polygenic and complex signal of selection in the face of multiple selective pressures, highlighting putatively adaptive variation between the six candidate CUs.

### Hierarchical population structure informs conservation units

As a first step toward delimiting conservation units for anadromous Arctic Char, we used a combination of common methods to assess the genetic population structure of our 30 sampling sites. This approach revealed a complex but geographically coherent and hierarchical picture of genetic structure.

At the broadest level, visible in the PCA (PC1) and NGSadmix (K = 2) results, the system is defined by an admixture cline from the Arctic lineage in the northwest to the Atlantic lineage in the southeast, as explored in detail in Dallaire et al. (2025). According to mitochondrial data, the divergence time between those lineages could reach up to 400,000 years, predating many glacial cycles (Jacobsen et al., 2022). Regardless of whether admixture occurred during previous interglacial periods, a prolonged phase of allopatry allowed these lineages to diverge significantly, which explains why much of the genetic variation observed in our current dataset reflects this secondary contact. This pattern provides an initial framework for delineating conservation units, as it reveals two distinct evolutionary trajectories: Northern sites are dominated by the Arctic lineage, while Southern sites exhibit varying degrees of admixture between the Arctic and Atlantic lineages. The distinctiveness between these two population segments is also supported by a low effective migration surface between the Kitikmeot and Kivalliq region, as well as Northern and Southern Baffin Island. Although the Northern and Southern populations do not represent entirely pure lineages, we argue that they should be managed as separate broad-scale conservation units. The persistence of this historical signal in contemporary patterns of variation suggests that secondary contact has not erased the imprint of prolonged allopatry. This dominance of ancient divergence over more recent demographic processes is particularly striking in the Arctic, where deglaciation occurred only a few thousand years ago (Dalton et al., 2020), and resembles findings in other post-glacial populations of fishes such as salmonids (Lehnert et al., 2019) and sticklebacks (Kirch et al., 2021), where deep lineage histories continue to structure modern diversity.

However, our analysis of structure revealed additional hierarchical levels of population structure within the North and South groups, where most individuals showed pure ancestry to a genetic cluster specific to its sampling site. While the ΔK summary statistic suggested 2 groups, studies have shown that this statistic is strongly biased toward K = 2 (Janes et al., 2017). In our study, this is not surprising given the overwhelming signal of lineage ancestry that this level of structure represents. Considering that ΔK was stable from K = 4 to 30 and that significant structure between rivers is expected according to the philopatric nature of Arctic Char reproductive migrations (Gyselman, 1994; Moore et al., 2013; Dallaire et al., 2021), we argue that all 30 clusters identified by NGSadmix could represent populations with some degree of demographic independence. These results highlight the strong hierarchical nature of genetic structure in Arctic Char, as observed in other anadromous salmonids (e.g. Beacham et al., 2004; Habicht et al., 2007; Dionne et al., 2008; Gilbert-Horvath et al., 2016; Östergren et al., 2021). Importantly, this fine-scale, river-specific structure aligns with current fisheries management practices, which often implement river-specific quotas and management measures to reflect the biological reality of Arctic Char populations.

Our sampling strategy focused on broad coverage across the Canadian range of anadromous Arctic Char rather than fine-scale resolution. As such, the six candidate CUs proposed here each cover coastal areas ranging for hundreds to thousands of km, which is much larger than current CUs in other salmonids in southern Canada (e.g., Xuereb et al., 2022; Lehnert et al., 2023). However, our analyses still uncovered patterns that may hold significance for local management. For example, we identified evidence of straying between most pairs of rivers inside 100 km from each other, and in one instance, up to 400 km (one fish sampled at EKA was genetically closer to fish in COP). Despite their philopatric behavior, Arctic Chars are known to show less fidelity to their natal site during the upstream migration in non-reproductive years (Gyselman, 1994; Klemetsen et al., 2003; Moore et al., 2013). When migrating strictly for overwintering purposes, fish might select watersheds with easily accessible winter habitats, for example the Ekalluk system in the Kitikmeot Region (EKA; Moore et al., 2017). In the case of AKL and CRB (Kivalliq), fish from either site were indistinguishable using whole-genome data, which suggests either panmixia between the two systems or that sampling at these sites is not targeting individuals during the upstream reproductive migration, but rather a mixed-stock of poorly differentiated source populations. Proper delimitation of MUs in Arctic Char would warrant additional local studies of genetic structure and migration, as was implemented in areas sustaining commercial and subsistence fisheries, such as in Cambridge Bay (Harris, Moore, et al., 2016; Moore et al., 2017), Cumberland Sound (Harris et al., 2014), Paulatuk (Harris, Boguski, et al., 2016), and Ulukhaktok (Lea et al., 2023). Going forward, whole-genome sequencing is likely not the most cost-effective method to tackle such local-scale questions. Alternatively, SNP panels can be developed to assign fish caught in mixed-stock fisheries to their population of origin, which holds promise for the accurate monitoring of fish stocks and the conciliation of commercial and subsistence fishing (Beemelmanns et al., 2025).

We chose K = 7 as an intermediate level of structure that could be used to delimit candidate CUs in Canadian anadromous Arctic Char. This level of structure subdivides our top-level Northern and Southern CUs, described earlier, and creates genetic groupings that conveniently, but somewhat unexpectedly, closely match the administrative regions of the Canadian Arctic (Fig. 1). A notable exception, however, is the union of both the Kivalliq (western) and Nunavik (eastern) coasts of the Hudson Bay in a single cluster (both coasts display signs of common ancestry until K = 17), while the rest of the Nunavik populations, around Ungava Bay, appeared as a distinct unit. Considering that K = 6 and K = 7 discriminated HRN and KJA (Fig. S4), our two westernmost sites, we suggest that these two could be paired to form a sixth CU, until additional sampling is performed in the Inuvialuit Settlement Region.

The choice of a level of genetic structure to define CUs implies subjectivity to some degree (Waples, 1995), as genetic differentiation typically occurs along a continuum rather than in clear-cut divisions. In the case of Arctic Char, many populations at the edge of candidate CUs display admixed ancestry to multiple clusters. However, the borders of the proposed CUs were supported by shifts in the dominant cluster that were concordant with areas of low effective migration in our EEMS analyses, as was for example observed in multiple marine species in Atlantic Canada (Wilcox et al., 2023). In the present study, candidate boundaries included the Boothia Peninsula between the Kitikmeot and the North Baffin CUs, and the Foxe Basin and Hudson Strait between the South Baffin and Hudson Bay/Ungava Bay CUs. While EEMS areas of low effective migration do not necessarily imply physical barriers to gene flow (Petkova et al., 2016), they should coincide with shifts in allele frequency that are relevant for informing conservation actions.

### Detecting local adaptation using whole-genome data

We further used our genomic dataset to detect signals of local adaptation in Arctic Char populations to support the delineation of CUs that also reflect patterns of local adaptation to divergent environments (sensu Funk et al., 2012). By using multiple analyses on millions of genetic markers, we obtained an overwhelmingly polygenic signal at the SNP-level, with numerous peaks of differentiation or associations with the environment across all linkage groups. To summarise this amount of noise and account for the high rate of false positives in the detection of adaptation, we 1) opted for a Windowed Z Analysis (WZA) to identify top candidate genes while accounting for linkage, and 2) focused on genes that were top candidates in multiple analyses.

As such, we were interested in the relationship between signals of local adaptation and known genomic regions of low recombination, as these might either be the result of hard selective sweeps or genomic features such as inversions or centromeres (Lotterhos, 2019). Our previous work inferred the position of many long haplotype blocks (i.e. sequences containing many polymorphisms inherited as a unit), suggesting a very low recombination rate within those segments (Dallaire et al. 2025). The haplotype frequencies of many of those putative haploblocks were heavily correlated with the admixture gradient between the Arctic and Atlantic glacial lineages. This implies that they act as local ancestry tracts and are much more constrained by post-glacial demography than spatial variation in selection, though the two are not mutually exclusive. In the partial RDA, we included each population’s average ancestry proportion as a covariate in the model, so that the analysis only tests for associations between genotypes and environment once the effect of lineage admixture has been factored out. Surprisingly, both putative local ancestry tracts and other local PCA outliers (identified in Dallaire et al. 2025) were enriched in top candidates from pRDA1, despite the partial RDA specifically controlling for the effect of Arctic-Atlantic ancestry.

However, RDA2 and pRDA1 identified common top candidates in a region of the LG13 identified as a potential inversion in Dallaire et al. (2025) (Fig 5a,c). This 290 kb region overlaps seven genes, including the insulin-like 5a and a protein regulating mitochondrial genes, hinting at their importance in metabolism or energy production. The alternative karyotype for this putative inversion is present in higher frequencies in both North and South Baffin, which present contrasted habitats compared to the rest of the Canadian Arctic, with fjord-like topography (watersheds with a higher slope and lower lake cover) and colder sea-surface temperatures. It is thus possible that this putative inversion could play a role in adaptation to these environments. Chromosomal inversions suppress recombination and are known to promote co-adaptation of neighbouring genes in the face of gene flow (Dobzhansky & Sturtevant, 1938; Kirkpatrick & Barton, 2006; Wellenreuther et al., 2019). This has previously been documented in rainbow trout (*Onchorhyncus mykiss*, Pearse et al., 2019), Atlantic Cod (*Gadus morhua*, Matschiner et al., 2022), Atlantic Herring (*Clupea harengus*, Jamsandekar et al., 2024), and Atlantic Silverside (*Menidia menidia*, Tigano et al., 2021). Another larger (1.2 Mb) candidate inversion on LG12 was found to be polymorphic in Arctic Char populations from Kitikmeot (Hale et al., 2021), but we did not find well-supported top candidate genes in this genomic region.

When using multiple genome scans and GEA methods with their respective biases, giving greater weight to markers identified by more than one method can help to reduce false positives (Forester et al., 2018). By doing so, we reduced our list of top candidates from 2,635 to 541 genes widely distributed across the genome. Despite low statistical support for GO enrichment after correction for multiple testing, we identified key gene functions related to lipid homeostasis and catabolic processes, which ranked among the best-supported GO categories across multiple sets of candidate genes. Lipid metabolism plays a critical role in the adaptation to Arctic climates, supporting membrane fluidity in cold-water ectotherms such as fishes (Ernst et al., 2016; Wang et al., 2021) and energy reserves during extreme dietary challenges such as prolonged fasting and hibernation (Olsen et al., 2021). The importance of fatty acid homeostasis in local adaptation was also observed in subarctic Atlantic Salmon populations, where genes regulating lipid metabolism were identified as drivers of adaptation (Lehnert et al., 2023).

In summary, identifying the biological processes and molecular functions involved in polygenic adaptation to multiple selective pressures remains challenging (Pritchard and Di Rienzo 2010; Le Corre and Kremer 2012; Rees et al. 2020). This is particularly true when using indirect inference of selection in the absence of phenotypic data on species with complex life cycles. Here, genome-wide, high-density SNP data provided unmatched resolution in the detection of signals of selection but required considerable data aggregation (e.g., through the WZA, as recently used in plants, Battlay et al., 2023; and *Drosophila*, Nunez et al., 2024) to make the results interpretable. This supported the importance of considering the relation between selection and recombination in our data, as highly linked regions of the genome could either be a consequence of selective sweeps or could have biased the detection of adaptation.

### Putatively adaptive variation reinforces candidate CUs

Despite challenges in detecting specific targets for selection, we used the overall putatively adaptive genomic variation to highlight groups of populations that could show local adaptation to similar habitats, thus supporting the candidate Conservation Units described earlier. For example, the distribution of allele frequencies in relation to environmental variation, as described by the pRDA (Fig. 4c-d), emphasizes one of our main findings—that a distinction based on water bodies in the Ungava and Hudson Bay groups was more significant than one based on landmasses (i.e., Nunavik vs. Kivalliq). These waterbodies mainly differed in coastal sea-surface temperature and salinity, and watersheds flowing in them are topographically distinct: Nunavik watersheds are generally less steep and have higher lake cover compared with those in South Baffin, our third recommended Conservation Unit in the South Arctic region.

In the North, environmental and geographical distances between populations were too confounded to support isolation-by-environment, but putatively adaptive genetic variation supported the distinction between the Kivalliq and North Baffin CUs, as well as the inclusion of the Abernathy Lake (ABL) population, east of the Boothia Peninsula, in the North Baffin CU. Similar to our results on genome-wide variation, we observed a weaker shift in allele frequencies associated with the environment between Kitikmeot and Inuvialuit Settlement Region populations, putting into question the distinction between those two candidate CUs. However, following a precautionary approach, we recommend maintaining these population segments as separate CUs, with boundaries that could be refined through additional sampling at the western limit of our study area.

Across our study area, conclusions based on genome-wide, neutral, and putatively adaptive genetic variation were largely consistent, with the adaptive data reinforcing the neutral patterns rather than providing novel insights. While one of the promises of population genomics was to reveal previously undetected adaptive groups that could be considered for conservation (Funk et al., 2012), there is growing evidence that genome-wide variation effectively predicts adaptive variation in many systems (Chhina et al., 2024). This was observed in other anadromous salmonids like Atlantic Salmon (Moore et al., 2014; Lehnert et al., 2023) and Coho Salmon (Xuereb et al., 2022), and this observation could be influenced by a few key factors. First, polygenic traits are expected to create weak selection signals (as identified in this study) that do not necessarily differ from neutral population structure (Pritchard et al., 2010). In contrast, large-effect loci such as GREB1L in steelhead (anadromous *O. mykiss*) and Chinook Salmon (*O. tshawytscha*), linked to migration timing, and vgll3 in Atlantic Salmon (*Salmo salar*), associated with age at maturity, underlie key phenotypes that are now explicitly considered in conservation and management (Waples et al., 2022).

Second, isolation-by-distance patterns suggest limited dispersal over large geographic scales (Aguillon et al., 2017) and straying is likely more frequent among watersheds sharing environmental characteristics. This could be either because of migratory cues such as water chemistry that differ among regional river systems (Dittman et al., 1996) or because of selection against maladapted migrants (Peterson et al., 2014) at local scales. There is in fact a long-standing hypothesis that homing evolved in anadromous salmonids as it promotes advantageous local adaptations (Quinn, 1993; McDowall, 2001; Keefer & Caudill, 2014). In contrast to the long marine migrations of other anadromous salmonids to offshore feeding areas, Arctic Char feed in coastal waters close to their natal rivers in summer (Dempson & Kristofferson, 1987; Spares et al., 2015; Moore et al., 2016), extending the potential for local adaptation to marine habitats in the species (Dallaire et al., 2021). As both freshwater and marine environments tend to be spatially autocorrelated (Legendre, 1993), it is not surprising that a significant part of the putatively adaptive genetic variation would be concordant with the neutral structure.

In this study, we reanalyzed genomic data from Arctic Char across the Canadian Arctic and provided a detailed assessment of population structure and connectivity. From this analysis, we identified six candidate Conservation Units (CUs) within two major groups displaying distinct evolutionary trajectories. This framework offers an important baseline for understanding intraspecific diversity at a national level, which can be further refined as additional data become available. Incorporating environmental, ecological, and life-history information alongside genomic insights will be essential for improving the resolution of these units and ensuring their alignment with meaningful biological boundaries. In that regard, Inuit ecological traditional knowledge will be invaluable as it includes detailed and fine-scale observations on distinct traits between population, e.g. size, taste and color, migratory behavior, and phenology.

Ultimately, this refined understanding of Arctic Char population structure can guide resource management strategies, ensuring the protection of locally adapted populations and their associated roles in the ecology of northern freshwater ecosystems, their importance for food security, and their contribution to the development of a sustainable economy in the Canadian Arctic. Future analyses that integrate genomic baselines with monitoring programs, harvest data, and indigenous traditional knowledge could improve the practical implementation of CUs. Such an approach would allow managers to better anticipate shifts in population dynamics, adjust harvest regulations proactively, and strengthen co-management frameworks with northern communities.

## Acknowledgement

This work was supported by a Large-Scale Applied Research Project grant from Genome Canada named ‘FISHES: Fostering Indigenous Small-scale Fisheries for Health, Economy and Food Security’. Sampling was made possible by the collaboration of Fisheries and Ocean Canada; Ministère de l’Environnement, de la Lutte contre les changements climatiques, de la Faune et des Parcs (Québec; Julien Mainguy); Taloyoak Umaruliririgut Association; Government of Nunavut (Zoya Martin); Makivik Corporation (Nunavik); and numerous other Inuit communities, organisations and local fishers across Canada. Many thanks to Bérénice Bougas, Charles Babin, Isabeau Caza-Allard, Louis-Philippe Collin, Alysse Perreault-Payette and Gabriel Piette-Lauzière for their help with laboratory work and coordination, as well as Raphaël Bouchard, Sann Delaive, Amanda Xuereb for their support and suggestions during the investigation of the data.

## Data Archiving Statement

All *Salvelinus alpinus* raw sequencing data analysed in this manuscript is available on Short Read Archive as part of project PRJNA1031558. A bioinformatical pipeline presenting the main lcWGS analyses is available at: https://github.com/xav9536/angsd_pipeline.

## Reference

Aguillon, S. M., Fitzpatrick, J. W., Bowman, R., Schoech, S. J., Clark, A. G., Coop, G., & Chen, N. (2017). Deconstructing isolation-by-distance: The genomic consequences of limited dispersal. PLoS Genetics, 13(8), 1–27. 10.1371/journal.pgen.1006911

Assis, J., Fernández Bejarano, S. J., Salazar, V. W., Schepers, L., Gouvêa, L., Fragkopoulou, E., Leclercq, F., Vanhoorne, B., Tyberghein, L., Serrão, E. A., Verbruggen, H., & De Clerck, O. (2024). Bio-ORACLE v3.0. Pushing marine data layers to the CMIP6 Earth System Models of climate change research. Global Ecology and Biogeography, 33(4), e13813. 10.1111/geb.13813

Avise, J. C., Arnold, J., Ball, R. M., Bermingham, E., Lamb, T., Neigel, J. E., Reeb, C. A., & Saunders, N. C. (1987). Intraspecific Phylogeography: The Mitochondrial DNA Bridge Between Population Genetics and Systematics. Annual Review of Ecology and Systematics, 18, 489–522.

Barnes, R., & Sahr, K. (2024). dggridR: Discrete Global Grids. https://CRAN.R-project.org/package=dggridR

Battlay, P., Wilson, J., Bieker, V. C., Lee, C., Prapas, D., Petersen, B., Craig, S., Van Boheemen, L., Scalone, R., De Silva, N. P., Sharma, A., Konstantinović, B., Nurkowski, K. A., Rieseberg, L. H., Connallon, T., Martin, M. D., & Hodgins, K. A. (2023). Large haploblocks underlie rapid adaptation in the invasive weed Ambrosia artemisiifolia. Nature Communications, 14(1), 1717. 10.1038/s41467-023-37303-4

Beacham, T. D., Lapointe, M., Candy, J. R., McIntosh, B., MacConnachie, C., Tabata, A., Kaukinen, K., Deng, L., Miller, K. M., & Withler, R. E. (2004). Stock Identification of Fraser River Sockeye Salmon Using Microsatellites and Major Histocompatibility Complex Variation. Transactions of the American Fisheries Society, 133(5), 1117–1137. 10.1577/T04-001.1

Beemelmanns, A., Bouchard, R., Michaelides, S., Normandeau, E., Jeon, H.-B., Chamlian, B., Babin, C., Hénault, P., Perrot, O., Harris, L. N., Zhu, X., Fraser, D., Bernatchez, L., & Moore, J.-S. (2025). Development of SNP Panels from Low-Coverage Whole Genome Sequencing (lcWGS) to Support Indigenous Fisheries for Three Salmonid Species in Northern Canada. Molecular Ecology Resources, e14040. 10.1111/1755-0998.14040

Benjamini, Y., & Hochberg, Y. (1995). Controlling the False Discovery Rate: A Practical and Powerful Approach to Multiple Testing. Journal of the Royal Statistical Society: Series B (Methodological), 57(1), 289–300. 10.1111/j.2517-6161.1995.tb02031.x

Booker, T. R., Yeaman, S., Whiting, J. R., & Whitlock, M. C. (2024). The WZA : A window based method for characterizing genotype–environment associations. Molecular Ecology Resources, 24(2), e13768. 10.1111/1755-0998.13768

Brubacher. (2004). An overview of Nunavut fisheries (p. 93) [Report]. Government of Nunavut.

Camacho, C., Coulouris, G., Avagyan, V., Ma, N., Papadopoulos, J., Bealer, K., & Madden, T. L. (2009). BLAST+: architecture and applications. BMC Bioinformatics, 10(1), 421. 10.1186/1471-2105-10-421

Chhina, A. K., Fernandez-Fournier, P., Lewthwaite, J., Booker, T., & Mooers, A. (2024). Data from: Does genome-wide variation and putatively adaptive variation identify the same set of distinct populations? (Version 4, p. 271770344 bytes) [Dataset]. Dryad. 10.5061/DRYAD.NVX0K6F1J

Christensen, K. A., Rondeau, E. B., Minkley, D. R., Leong, J. S., Nugent, C. M., Danzmann, R. G., Ferguson, M. M., Stadnik, A., Devlin, R. H., Muzzerall, R., Edwards, M., Davidson, W. S., & Koop, B. F. (2021). Retraction: The Arctic charr (Salvelinus alpinus) genome and transcriptome assembly. PLOS ONE, 16(2), e0247083. 10.1371/journal.pone.0247083

Coates, D. J., Byrne, M., & Moritz, C. (2018). Genetic Diversity and Conservation Units: Dealing With the Species-Population Continuum in the Age of Genomics. Frontiers in Ecology and Evolution, 6. https://www.frontiersin.org/articles/10.3389/fevo.2018.00165

COSEWIC. (2010). COSEWIC assessment and status report on the Atlantic Salmon Salmo salar (Nunavik population, Labrador population, Northeast Newfoundland population, South Newfoundland population, Southwest Newfoundland population, Northwest Newfoundland population, Quebec Eastern North Shore population, Quebec Western North Shore population, Anticosti Island population, Inner St. Lawrence population, Lake Ontario population, Gaspé-Southern Gulf of St. Lawrence population, Eastern Cape Breton population, Nova Scotia Southern Upland population, Inner Bay of Fundy population, Outer Bay of Fundy population) in Canada (p. xlvii + 136). Committee on the Status of Endangered Wildlife in Canada.

Dallaire, X., Bouchard, R., Hénault, P., Ulmo-Diaz, G., Normandeau, E., Mérot, C., Bernatchez, L., & Moore, J.-S. (2023). Widespread Deviant Patterns of Heterozygosity in Whole-Genome Sequencing Due to Autopolyploidy, Repeated Elements, and Duplication. Genome Biology and Evolution, 15(12), evad229. 10.1093/gbe/evad229

Dallaire, X., Normandeau, E., Brazier, T., Harris, L., Michael, M. M. H., Mérot, C., & Moore, J.-S. (2025). Leveraging whole genomes, mitochondrial DNA, and haploblocks to decipher complex demographic histories: an example from a broadly admixed arctic fish. Molecular Ecology. 10.1101/2024.12.11.628006

Dallaire, X., Normandeau, É., Mainguy, J., Tremblay, J.-É., Bernatchez, L., & Moore, J.-S. (2021). Genomic data support management of anadromous Arctic Char fisheries in Nunavik by highlighting neutral and putatively adaptive genetic variation. Evolutionary Applications, n/a(n/a). 10.1111/eva.13248

Dalton, A. S., Margold, M., Stokes, C. R., Tarasov, L., Dyke, A. S., Adams, R. S., Allard, S., Arends, H. E., Atkinson, N., Attig, J. W., Barnett, P. J., Barnett, R. L., Batterson, M., Bernatchez, P., Borns, H. W., Breckenridge, A., Briner, J. P., Brouard, E., Campbell, J. E., … Wright, H. E. (2020). An updated radiocarbon-based ice margin chronology for the last deglaciation of the North American Ice Sheet Complex. Quaternary Science Reviews, 234. 10.1016/j.quascirev.2020.106223

Dauphin, B., Rellstab, C., Wüest, R. O., Karger, D. N., Holderegger, R., Gugerli, F., & Manel, S. (2023). Re-thinking the environment in landscape genomics. Trends in Ecology & Evolution, 38(3), 261–274. 10.1016/j.tree.2022.10.010

Dempson, J. B., & Kristofferson, A. H. (1987). Spatial and Temporal Aspects of the Ocean Migration of Anadromous Arctic Char. American Fisheries Society Symposium, 1, 340–357.

DFO. (2024). Pacific Salmon Outlook: Pacific Region. Fisheries and Oceans Canada. https://waves-vagues.dfo-mpo.gc.ca/library-bibliotheque/41263625.pdf

Dionne, M., Caron, F., Dodson, J. J., & Bernatchez, L. (2008). Landscape genetics and hierarchical genetic structure in Atlantic salmon: The interaction of gene flow and local adaptation. Molecular Ecology, 17(10), 2382–2396. 10.1111/j.1365-294X.2008.03771.x

Dittman, A. H., Quinn, T. P., & Nevitt, G. A. (1996). Timing of imprinting to natural and artificial odors by coho salmon (*Oncorhynchus kisutch*). Canadian Journal of Fisheries and Aquatic Sciences, 53(2), 434–442. 10.1139/f95-185

Dobzhansky, T., & Sturtevant, A. H. (1938). Inversions in the Chromosomes of Drosophila Pseudoobscura. Genetics, 23(1), 28–64.

Dubos, V., St-Hilaire, A., & Bergeron, N. E. (2023). Fuzzy logic modelling of anadromous Arctic char spawning habitat from Nunavik Inuit knowledge. Ecological Modelling, 477, 110262. 10.1016/j.ecolmodel.2022.110262

Ellegren, H. (2004). Microsatellites: simple sequences with complex evolution. Nature Reviews Genetics, 5(6), 435–445. 10.1038/nrg1348

Ernst, R., Ejsing, C. S., & Antonny, B. (2016). Homeoviscous Adaptation and the Regulation of Membrane Lipids. Journal of Molecular Biology, 428(24, Part A), 4776–4791. 10.1016/j.jmb.2016.08.013

Evanno, G., Regnaut, S., & Goudet, J. (2005). Detecting the number of clusters of individuals using the software STRUCTURE : a simulation study. Molecular Ecology, 14(8), 2611–2620. 10.1111/j.1365-294X.2005.02553.x

Forest, F., Grenyer, R., Rouget, M., Davies, T. J., Cowling, R. M., Faith, D. P., Balmford, A., Manning, J. C., Procheş, Ş., van der Bank, M., Reeves, G., Hedderson, T. A. J., & Savolainen, V. (2007). Preserving the evolutionary potential of floras in biodiversity hotspots. Nature, 445(7129), 757–760. 10.1038/nature05587

Forester, B. R., Lasky, J. R., Wagner, H. H., & Urban, D. L. (2018). Comparing methods for detecting multilocus adaptation with multivariate genotype– environment associations. Molecular Ecology, 27(9), 2215–2233. 10.1111/mec.14584

Fox, E. A., Wright, A. E., Fumagalli, M., & Vieira, F. G. (2019). ngsLD: evaluating linkage disequilibrium using genotype likelihoods. Bioinformatics, 35(19), 3855–3856. 10.1093/bioinformatics/btz200

Frankham, R., Ballou, J. D., Dudash, M. R., Eldridge, M. D. B., Fenster, C. B., Lacy, R. C., Mendelson, J. R., Porton, I. J., Ralls, K., & Ryder, O. A. (2012). Implications of different species concepts for conserving biodiversity. Biological Conservation, 153, 25–31. 10.1016/j.biocon.2012.04.034

Fraser, D. J., & Bernatchez, L. (2001). Adaptive evolutionary conservation: Towards a unified concept for defining conservation units. Molecular Ecology, 10(12), 2741–2752. 10.1046/j.1365-294X.2001.t01-1-01411.x

Friesen, T. M. (2004). Contemporaneity of Dorset and Thule Cultures in the North American Arctic: New Radiocarbon Dates from Victoria Island, Nunavut. Current Anthropology, 45(5), 685–691. 10.1086/425635

Funk, W. C., McKay, J. K., Hohenlohe, P. A., & Allendorf, F. W. (2012). Harnessing genomics for delineating conservation units. Trends in Ecology and Evolution, 27(9), 489–496. 10.1016/j.tree.2012.05.012

Garnett, S. T., & Christidis, L. (2017). Taxonomy anarchy hampers conservation. Nature, 546(7656), 25–27. 10.1038/546025a

Gascuel, O. (1997). BIONJ: an improved version of the NJ algorithm based on a simple model of sequence data. Molecular Biology and Evolution, 14(7), 685–695. 10.1093/oxfordjournals.molbev.a025808

Gautier, M. (2015). Genome-wide scan for adaptive divergence and association with population-specific covariates. Genetics, 201(4), 1555–1579. 10.1534/genetics.115.181453

Gilbert-Horvath, E. A., Pipal, K. A., Spence, B. C., Williams, T. H., & Garza, J. C. (2016). Hierarchical Phylogeographic Structure of Coho Salmon in California. Transactions of the American Fisheries Society, 145(5), 1122–1138. 10.1080/00028487.2016.1201003

Green, D. M. (2005). Designatable Units for Status Assessment of Endangered Species. Conservation Biology, 19(6), 1813–1820. 10.1111/j.1523-1739.2005.00284.x

Gyselman, E. C. (1994). Fidelity of Anadromous Arctic Char. Canadian Journal of Fisheries and Aquatic Sciences, 51, 1927–1934.

Habicht, C., Seeb, L. W., & Seeb, J. E. (2007). Genetic and Ecological Divergence Defines Population Structure of Sockeye Salmon Populations Returning to Bristol Bay, Alaska, and Provides a Tool for Admixture Analysis. Transactions of the American Fisheries Society, 136(1), 82–94. 10.1577/T06-001.1

Hale, M. C., Campbell, M. A., & McKinney, G. J. (2021). A candidate chromosome inversion in Arctic charr (*Salvelinus alpinus*) identified by population genetic analysis techniques. G3 Genes|Genomes|Genetics, jkab267. 10.1093/g3journal/jkab267

Harris, L. N., Boguski, D. A., Gallagher, C. P., & Howland, K. L. (2016). Genetic Stock Identification and Relative Contribution of Arctic Char (*Salvelinus alpinus*) from the Hornaday and Brock Rivers to Subsistence Fisheries in Darnley Bay, Northwest Territories + Supplementary Appendix Tables S1 to S4 (See Article Tools). ARCTIC, 69(3), 231–245. 10.14430/arctic4578

Harris, L. N., Moore, J.-S., Bajno, R., & Tallman, R. F. (2016). Genetic Stock Structure of Anadromous Arctic Char in Canada’s Central Arctic: Potential Implications for the Management of Canada’s Largest Arctic Char Commercial Fishery. North American Journal of Fisheries Management, 36(6), 1473–1488. 10.1080/02755947.2016.1227399

Harris, L. N., Moore, J.-S., Galpern, P., Tallman, R. F., & Taylor, E. B. (2014). Geographic influences on fine-scale, hierarchical population structure in northern Canadian populations of anadromous Arctic Char (Salvelinus alpinus). Environmental Biology of Fishes, 97(11), 1233–1252. 10.1007/s10641-013-0210-y

Harris, L. N., Yurkowski, D. J., Gilbert, M. J. H., Else, B., Duke, P. J., Ahmed, M., Tallman, R. F., Fisk, A. T., & Moore, J. S. (2020). Depth and temperature preference of anadromous Arctic Char, Salvelinus alpinus, in the Kitikmeot Sea: a shallow and low salinity area of the Canadian Arctic. Marine Ecology Progress Series, 634, 175–197. 10.3354/meps13195

Jacobsen, M. W., Jensen, N. W., Nygaard, R., Præbel, K., Jónsson, B., Nielsen, N. H., Pujolar, J. M., Fraser, D. J., Bernatchez, L., & Hansen, M. M. (2022). A melting pot in the Arctic: Analysis of mitogenome variation in Arctic char (*Salvelinus alpinus*) reveals a 1000 km contact zone between highly divergent lineages. Ecology of Freshwater Fish, 31(2), 330–346. 10.1111/eff.12633

Jamsandekar, M., Ferreira, M. S., Pettersson, M. E., Farrell, E. D., Davis, B. W., & Andersson, L. (2024). The origin and maintenance of supergenes contributing to ecological adaptation in Atlantic herring. Nature Communications, 15(1), 9136. 10.1038/s41467-024-53079-7

Janes, J. K., Miller, J. M., Dupuis, J. R., Malenfant, R. M., Gorrell, J. C., Cullingham, C. I., & Andrew, R. L. (2017). The K = 2 conundrum. Molecular Ecology, 26(14), 3594–3602. 10.1111/mec.14187

Keefer, M. L., & Caudill, C. C. (2014). Homing and straying by anadromous salmonids: A review of mechanisms and rates. Reviews in Fish Biology and Fisheries, 24(1), 333–368. 10.1007/s11160-013-9334-6

Kess, T., Dempson, J. B., Lehnert, S. J., Layton, K., Einfeldt, A., Bentzen, P., Salisbury, S. J., Messmer, A. M., Duffy, S., Ruzzante, D. E., Nugent, C. M., Ferguson, M. M., Leong, J. S., Koop, B. F., O’Connell, M. F., & Bradbury, I. R. (2021). Genomic basis of deep water adaptation in Arctic Charr (*Salvelinus alpinus*) morphs. Molecular Ecology, mec.16033. 10.1111/mec.16033

Kirch, M., Romundset, A., Gilbert, M. T. P., Jones, F. C., & Foote, A. D. (2021). Ancient and modern stickleback genomes reveal the demographic constraints on adaptation. Current Biology, 31(9), 2027–2036.e8. 10.1016/j.cub.2021.02.027

Kirkpatrick, M., & Barton, N. (2006). Chromosome inversions, local adaptation and speciation. Genetics, 173(1), 419–434. 10.1534/genetics.105.047985

Klemetsen, A., Amundsen, P.-A., Dempson, J. B., Jonsson, B., Jonsson, N., O’Connell, M. F., & Mortensen, E. (2003). Atlantic salmon Salmo salar L., brown trout Salmo trutta L. and Arctic charr Salvelinus alpinus (L.): a review of aspects of their life histories. Ecology of Freshwater Fish, 12(1), 1–59. 10.1034/j.1600-0633.2003.00010.x

Klopfenstein, D. V., Zhang, L., Pedersen, B. S., Ramírez, F., Warwick Vesztrocy, A., Naldi, A., Mungall, C. J., Yunes, J. M., Botvinnik, O., Weigel, M., Dampier, W., Dessimoz, C., Flick, P., & Tang, H. (2018). GOATOOLS: A Python library for Gene Ontology analyses. Scientific Reports, 8, 10872. 10.1038/s41598-018-28948-z

Kopelman, N. M., Mayzel, J., Jakobsson, M., Rosenberg, N. A., & Mayrose, I. (2015). Clumpak: a program for identifying clustering modes and packaging population structure inferences across K. Molecular Ecology Resources, 15(5), 1179–1191. 10.1111/1755-0998.12387

Laikre, L., Hoban, S., Bruford, M. W., Segelbacher, G., Allendorf, F. W., Gajardo, G., Rodríguez, A. G., Hedrick, P. W., Heuertz, M., Hohenlohe, P. A., Jaffé, R., Johannesson, K., Liggins, L., MacDonald, A. J., OrozcoterWengel, P., Reusch, T. B. H., Rodríguez-Correa, H., Russo, I.-R. M., Ryman, N., & Vernesi, C. (2020). Post-2020 goals overlook genetic diversity. Science, 367(6482), 1083–1085. 10.1126/science.abb2748

Layton, K. K. S., Snelgrove, P. V. R., Dempson, J. B., Kess, T., Lehnert, S. J., Bentzen, P., Duffy, S. J., Messmer, A. M., Stanley, R. R. E., DiBacco, C., Salisbury, S. J., Ruzzante, D. E., Nugent, C. M., Ferguson, M. M., Leong, J. S., Koop, B. F., & Bradbury, I. R. (2021). Genomic evidence of past and future climate-linked loss in a migratory Arctic fish. Nature Climate Change, 11(2), 158–165. 10.1038/s41558-020-00959-7

Lea, E. V., Olokhaktomiut Hunters and Trappers Committee, & Harwood. (2023). Fish and Marine Mammals Harvested near Ulukhaktok, Northwest Territories, with a focus on Anadromous Arctic Char (Salvelinus alpinus) (No. 2023/014; p. 23). Fisheries and Oceans Canada.

Legendre, P. (1993). Spatial Autocorrelation: Trouble or New Paradigm? Ecology, 74(6), 1659–1673. 10.2307/1939924

Lehnert, S. J., Bentzen, P., Kess, T., Lien, S., Horne, J. B., Clément, M., & Bradbury, I. R. (2019). Chromosome polymorphisms track trans-Atlantic divergence and secondary contact in Atlantic salmon. Molecular Ecology, 28(8), 2074–2087. 10.1111/mec.15065

Lehnert, S. J., Bradbury, I. R., Wringe, B. F., Van Wyngaarden, M., & Bentzen, P. (2023). Multifaceted framework for defining conservation units: An example from Atlantic salmon (Salmo salar) in Canada. Evolutionary Applications, 16(9), 1568–1585. 10.1111/eva.13587

Li, P., van Coeverden de Groot, P., Clemente-Carvalho, R. B. G., & Lougheed, S. C. (2021). ddRAD genotyping reveals hierarchical genetic population structure in anadromous Arctic char (Salvelinus alpinus) in the Lower Northwest Passage, Nunavut. Canadian Journal of Fisheries and Aquatic Sciences, 78(4), 457–471. 10.1139/cjfas-2020-0069

Linderoth, T. (2018). Identifying Population Histories, Adaptive Genes, and Genetic Duplication from Population-Scale Next Generation Sequencing. University of California, Berkeley.

Lotterhos, K. E. (2019). The Effect of Neutral Recombination Variation on Genome Scans for Selection. G3 Genes|Genomes|Genetics, 9(6), 1851–1867. 10.1534/g3.119.400088

Luu, K., Bazin, E., & Blum, M. G. B. (2017). pcadapt: an R package to perform genome scans for selection based on principal component analysis. Molecular Ecology Resources, 17(1), 67–77. 10.1111/1755-0998.12592

Mace, G. M. (2004). The role of taxonomy in species conservation. Philosophical Transactions of the Royal Society of London. Series B: Biological Sciences, 359(1444), 711–719. 10.1098/rstb.2003.1454

Madsen, R. P. A., Jacobsen, M. W., O’Malley, K. G., Nygaard, R., Præbel, K., Jónsson, B., Pujolar, J. M., Fraser, D. J., Bernatchez, L., & Hansen, M. M. (2019). Genetic population structure and variation at phenology-related loci in anadromous Arctic char (Salvelinus alpinus). Ecology of Freshwater Fish, 00, 1–14. 10.1111/eff.12504

Marcus, J., Ha, W., Barber, R. F., & Novembre, J. (2021). Fast and flexible estimation of effective migration surfaces. eLife, 10, e61927. 10.7554/eLife.61927

Matschiner, M., Barth, J. M. I., Tørresen, O. K., Star, B., Baalsrud, H. T., Brieuc, M. S. O., Pampoulie, C., Bradbury, I., Jakobsen, K. S., & Jentoft, S. (2022). Supergene origin and maintenance in Atlantic cod. Nature Ecology & Evolution, 6(4), 469–481. 10.1038/s41559-022-01661-x

McDowall, R. M. (2001). Anadromy and homing: two life history traits with adaptive synergies in salmonid fishes? Fish and Fisheries, 2(1), 78–85. 10.1046/j.1467-2979.2001.00036.x

Meisner, J., & Albrechtsen, A. (2018). Inferring Population Structure and Admixture Proportions in Low-Depth NGS Data. Genetics, 210(2), 719–731. 10.1534/genetics.118.301336

Moore, J.-S., Bajno, R., Reist, J. D., & Taylor, E. B. (2015). Post-glacial recolonization of the North American Arctic by Arctic char (Salvelinus alpinus): Genetic evidence of multiple northern refugia and hybridization between glacial lineages. Journal of Biogeography, 42(11), 2089–2100. 10.1111/jbi.12600

Moore, J.-S., Bourret, V., Dionne, L., Bradbury, I. A. N., Reilly, P. O., & Kent, M. (2014). Conservation genomics of anadromous Atlantic salmon across its North American range : outlier loci identify the same patterns of population structure as neutral loci. Molecular Ecology, 23, 5680–5697. 10.1111/mec.12972

Moore, J.-S., Harris, L. N., Kessel, S. T., Bernatchez, L., Tallman, R. F., & Fisk, A. T. (2016). Preference for near-shore and estuarine habitats in anadromous Arctic char (Salvelinus alpinus) from the Canadian high Arctic (Victoria Island, NU) revealed by acoustic telemetry. Canadian Journal of Fisheries and Aquatic Sciences, 53(9), 1689–1699. 10.1017/CBO9781107415324.004

Moore, J.-S., Harris, L. N., Luyer, J. L., Sutherland, B., Rougemont, Q., Tallman, R. F., Fisk, A. T., & Bernatchez, L. (2017). Genomics and telemetry suggest a role for migration harshness in determining overwintering habitat choice, but not gene flow, in anadromous Arctic Char. Molecular Ecology, 1–14. 10.1111/mec.14393

Moore, J.-S., Harris, L. N., Tallman, R. F., & B, T. E. (2013). The interplay between dispersal and gene flow in anadromous Arctic char (Salvelinus alpinus): implications for potential for local adaptation. Canadian Journal of Fisheries and Aquatic Sciences, 1338(July), 1327–1338.

Moritz, C. (2002). Strategies to Protect Biological Diversity and the Evolutionary Processes That Sustain It. Systematic Biology, 51(2), 238–254. 10.1080/10635150252899752

Nunez, J. C. B., Lenhart, B. A., Bangerter, A., Murray, C. S., Mazzeo, G. R., Yu, Y., Nystrom, T. L., Tern, C., Erickson, P. A., & Bergland, A. O. (2024). A cosmopolitan inversion facilitates seasonal adaptation in overwintering *Drosophila*. GENETICS, 226(2), iyad207. 10.1093/genetics/iyad207

Oksanen, J., Simpson, G. L., Blanchet, F. G., Kindt, R., Legendre, P., Minchin, P. R., O’Hara, R. B., Solymos, P., Stevens, M. H. H., Szoecs, E., Wagner, H., Barbour, M., Bedward, M., Bolker, B., Borcard, D., Carvalho, G., Chirico, M., Caceres, M. D., Durand, S., … Weedon, J. (2022). vegan: Community Ecology Package. https://CRAN.R-project.org/package=vegan

Olsen, L., Thum, E., & Rohner, N. (2021). Lipid metabolism in adaptation to extreme nutritional challenges. Developmental Cell, 56(10), 1417–1429. 10.1016/j.devcel.2021.02.024

Östergren, J., Palm, S., Gilbey, J., Spong, G., Dannewitz, J., Königsson, H., Persson, J., & Vasemägi, A. (2021). A century of genetic homogenization in Baltic salmon—evidence from archival DNA. Proceedings of the Royal Society B: Biological Sciences, 288(1949), rspb.2020.3147, 20203147. 10.1098/rspb.2020.3147

Paradis, E., & Schliep, K. (2019). ape 5.0: an environment for modern phylogenetics and evolutionary analyses in R. Bioinformatics, 35(3), 526–528. 10.1093/bioinformatics/bty633

Pearse, D. E., Barson, N. J., Nome, T., Gao, G., Campbell, M. A., Abadía-Cardoso, A., Anderson, E. C., Rundio, D. E., Williams, T. H., Naish, K. A., Moen, T., Liu, S., Kent, M., Moser, M., Minkley, D. R., Rondeau, E. B., Brieuc, M. S. O., Sandve, S. R., Miller, M. R., … Lien, S. (2019). Sex-dependent dominance maintains migration supergene in rainbow trout. Nature Ecology & Evolution, 3(12), 1731–1742. 10.1038/s41559-019-1044-6

Peterson, D. A., Hilborn, R., & Hauser, L. (2014). Local adaptation limits lifetime reproductive success of dispersers in a wild salmon metapopulation. Nature Communications, 5(1), 3696. 10.1038/ncomms4696

Petkova, D., Novembre, J., & Stephens, M. (2016). Visualizing spatial population structure with estimated effective migration surfaces. Nature Genetics, 48(1), 94–100. 10.1038/ng.3464

Priest, H., & Usher, P. J. (2004). Nunavut Wildlife Harvest Study (p. 822). Nunavut Wildlife Management Board.

Pritchard, J. K., Pickrell, J. K., & Coop, G. (2010). The Genetics of Human Adaptation: Hard Sweeps, Soft Sweeps, and Polygenic Adaptation. Current Biology, 20(4), R208–R215. 10.1016/j.cub.2009.11.055

Quinn, T. P. (1993). A review of homing and straying of wild and hatchery-produced salmon. Fisheries Research, 18(1–2), 29–44. 10.1016/0165-7836(93)90038-9

Reist, J. D., Power, M., & Dempson, J. B. (2013). Arctic charr (Salvelinus alpinus): a case study of the importance of understanding biodiversity and taxonomic issues in northern fishes. Biodiversity, 14(1), 45–56. 10.1080/14888386.2012.725338

Rellstab, C., Gugerli, F., Eckert, A. J., Hancock, A. M., & Holderegger, R. (2015). A practical guide to environmental association analysis in landscape genomics. Molecular Ecology, 24(17), 4348–4370. 10.1111/mec.13322

Rosenblum, E. B., Sarver, B. A. J., Brown, J. W., Des Roches, S., Hardwick, K. M., Hether, T. D., Eastman, J. M., Pennell, M. W., & Harmon, L. J. (2012). Goldilocks Meets Santa Rosalia: An Ephemeral Speciation Model Explains Patterns of Diversification Across Time Scales. Evolutionary Biology, 39(2), 255–261. 10.1007/s11692-012-9171-x

Roux, C., Fraïsse, C., Romiguier, J., Anciaux, Y., Galtier, N., & Bierne, N. (2016). Shedding Light on the Grey Zone of Speciation along a Continuum of Genomic Divergence. PLOS Biology, 14(12), e2000234. 10.1371/journal.pbio.2000234

Roux, M.-J., Tallman, R. F., & Martin, Z. A. (2019). Small-scale fisheries in Canada’s Arctic: Combining science and fishers knowledge towards sustainable management. Marine Policy, 101, 177–186. 10.1016/j.marpol.2018.01.016

Ryder, O. (1986). Conservation and systematic: The dilemma of subspecies. Trends Ecol Evol, 1, 9–10.

Salisbury, S. J., McCracken, G. R., Keefe, D., Perry, R., & Ruzzante, D. E. (2019). Extensive secondary contact among three glacial lineages of Arctic Char (Salvelinus alpinus) in Labrador and Newfoundland. Ecology and Evolution, 9(4), 2031–2045. 10.1002/ece3.4893

Schindler, D. E., Hilborn, R., Chasco, B., Boatright, C. P., Quinn, T. P., Rogers, L. A., & Webster, M. S. (2010). Population diversity and the portfolio effect in an exploited species. Nature, 465(7298), 609–612. 10.1038/nature09060

Seehausen, O., Takimoto, G., Roy, D., & Jokela, J. (2008). Speciation reversal and biodiversity dynamics with hybridization in changing environments. Molecular Ecology, 17(1), 30–44. 10.1111/j.1365-294X.2007.03529.x

Skotte, L., Korneliussen, T. S., & Albrechtsen, A. (2013). Estimating Individual Admixture Proportions from Next Generation Sequencing Data. Genetics, 195(3), 693–702. 10.1534/genetics.113.154138

Spares, A. D., Stokesbury, M. J. W., Dadswell, M. J., O’Dor, R. K., & Dick, T. A. (2015). Residency and movement patterns of Arctic charr Salvelinus alpinus relative to major estuaries. Journal of Fish Biology, 86(6), 1754–1780. 10.1111/jfb.12683

Tallman, R. F., Roux, M.-J., & Martin, Z. A. (2019). Governance and assessment of small-scale data-limited Arctic Charr fisheries using productivity-susceptibility analysis coupled with life history invariant models. Marine Policy, 101, 187–197. 10.1016/j.marpol.2017.11.032

Therkildsen, N. O., & Palumbi, S. R. (2017). Practical low-coverage genomewide sequencing of hundreds of individually barcoded samples for population and evolutionary genomics in nonmodel species. Molecular Ecology Resources, 17(2), 194–208. 10.1111/1755-0998.12593

Tigano, A., Jacobs, A., Wilder, A. P., Nand, A., Zhan, Y., Dekker, J., & Therkildsen, N. O. (2021). Chromosome-Level Assembly of the Atlantic Silverside Genome Reveals Extreme Levels of Sequence Diversity and Structural Genetic Variation. Genome Biology and Evolution, 13(6), evab098. 10.1093/gbe/evab098

UniProt Consortium. (2024). Swiss-Prot: A manually annotated and reviewed protein sequence database. (Version 1.2) [Dataset].

Vieira, F. G., Lassalle, F., Korneliussen, T. S., & Fumagalli, M. (2016). Improving the estimation of genetic distances from Next-Generation Sequencing data. Biological Journal of the Linnean Society, 117(1), 139–149. 10.1111/bij.12511

Wang, C., Gong, Y., Deng, F., Ding, E., Tang, J., Codling, G., Challis, J. K., Green, D., Wang, J., Chen, Q., Xie, Y., Su, S., Yang, Z., Raine, J., Jones, P. D., Tang, S., & Giesy, J. P. (2021). Remodeling of Arctic char (*Salvelinus alpinus*) lipidome under a stimulated scenario of Arctic warming. Global Change Biology, 27(14), 3282–3298. 10.1111/gcb.15638

Waples, R. S. (1991). Pacific salmon, Oncorhynchus spp., and the definition of “species” under the Endangered Species Act. Marine Fisheries Review, 53(3), 11–22.

Waples, R. S. (1995). Evolutionarily significant units and the conservation of biological diversity under the Endangered Species Act. American Fisheries Society Symposium, 17, 8–27.

Waples, R. S., Ford, M. J., Nichols, K., Kardos, M., Myers, J., Thompson, T. Q., Anderson, E. C., Koch, I. J., McKinney, G., Miller, M. R., Naish, K., Narum, S. R., O’Malley, K. G., Pearse, D. E., Pess, G. R., Quinn, T. P., Seamons, T. R., Spidle, A., Warheit, K. I., & Willis, S. C. (2022). Implications of Large-Effect Loci for Conservation: A Review and Case Study with Pacific Salmon. Journal of Heredity, 113(2), 121–144. 10.1093/jhered/esab069

Wellenreuther, M., Mérot, C., Berdan, E., & Bernatchez, L. (2019). Going beyond SNPs: The role of structural genomic variants in adaptive evolution and species diversification. Molecular Ecology, 28(6), 1203–1209. 10.1111/mec.15066

Wilcox, M. A., Jeffery, N. W., DiBacco, C., Bradbury, I. R., Lowen, B., Wang, Z., Beiko, R. G., & Stanley, R. R. E. (2023). Integrating seascape resistances and gene flow to produce area-based metrics of functional connectivity for marine conservation planning. Landscape Ecology, 38(9), 2189–2205. 10.1007/s10980-023-01690-2

Xu, S., Dai, Z., Guo, P., Fu, X., Liu, S., Zhou, L., Tang, W., Feng, T., Chen, M., Zhan, L., Wu, T., Hu, E., Jiang, Y., Bo, X., & Yu, G. (2021). ggtreeExtra: Compact Visualization of Richly Annotated Phylogenetic Data. Molecular Biology and Evolution, 38(9), 4039–4042. 10.1093/molbev/msab166

Xuereb, A., Rougemont, Q., Dallaire, X., Moore, J.-S., Normandeau, E., Bougas, B., Perreault-Payette, A., Koop, B. F., Withler, R., Beacham, T., & Bernatchez, L. (2022). Re-evaluating Coho salmon (Oncorhynchus kisutch) conservation units in Canada using genomic data. Evolutionary Applications, 15(11), 1925–1944. 10.1111/eva.13489

Yu, G., Smith, D. K., Zhu, H., Guan, Y., & Lam, T. T.-Y. (2017). ggtree: an r package for visualization and annotation of phylogenetic trees with their covariates and other associated data. Methods in Ecology and Evolution, 8(1), 28–36. 10.1111/2041-210X.12628

