## Supplementary material for "Conservation units for anadromous Arctic Char (*Salvelinus alpinus*) in the Canadian Arctic informed by genetic structure, population connectivity and adaptive genomic variation": Fig. S

### Supplementary Tables

Table S1: Summary of environmental variables used for isolation-by-environment and Gene-Environment Associations analyses. Collinearity with other environmental variable and glacial lineage ancestry was estimated using Pearson’s r. Note that Precipitation was noted included in analyses, as it was strongly correlated to Air T°.

Table S2: Values for environmental variable, averaged over the catchment area for freshwater variables (green), and over a coastal zone around the sampled river mouth for marine variables (blue).

Table S3: Genomic regions identified as outlier after multidimensional scaling (MDS) on local principal component analyses (PCA) for windows of 100 SNPs in Dallaire et al. (2025). Local PCA outlier regions forming three distinct clusters which frequencies in populations correlate to average glacial lineage ancestry were marked as putative local ancestry tracts.

Table S4: Pairwise fixation index (F_ST_, below the diagonal) and allele frequency difference (AFD, above the diagonal) between all 30 populations.

Table S5: Mantel’s r and associated p-values estimated over 999 iterations for Mantel and partial Mantel tests for the relation between genetic distance (AFD or F_ST_) and marine (km, geographical distance between sampling sites), environmental (Euclidian distance from a principal component analysis of 11 environmental variables), or ancestry (difference in average ancestry for K = 2 in NGSadmix) distance matrices in all or either Northern or Southern populations. Environmental distance was further decomposed in individual variables (difference in value between populations) to test the effect of components on isolation-by-environment. Tests with p-values under 0.01 were highlighted in bold and gray.

Table S6: Number of top candidate genes by genome scan and Gene-Environment Association test. The number and proportion of those genes overlapping with putative local ancestry tracts and other local PCA outliers from Dallaire et al. (2025) were calculated, and the proportion was compared to the proportion of all genes with at least 5 SNPs overlapping those regions with χ^2^. Proportions in bold and gray are methods where putative local ancestry tracts and other local PCA outliers are enriched in top candidate genes.

Table S7: List of top candidate genes (empirical p-value < 0.001 in a windowed Z analysis) identified by either genome scan or Gene-Environment Association methods. Genes marked as top candidates by multiple methods are highlighted in gray, and overlapping local PCA outlier regions (Table S3, see Dallaire et al. 2025) are noted.

Table S8: Gene ontology (GO) terms enriched in top candidate gene lists from each genome scan or Gene-Environment Association method. GO terms with adjusted p-values under 0.05 are in bold.

### Supplementary Figures

Figure S1: Principal component analysis of environmental variation for 12 freshwater and marine variables. Sampling sites are colored according to their geographical region.

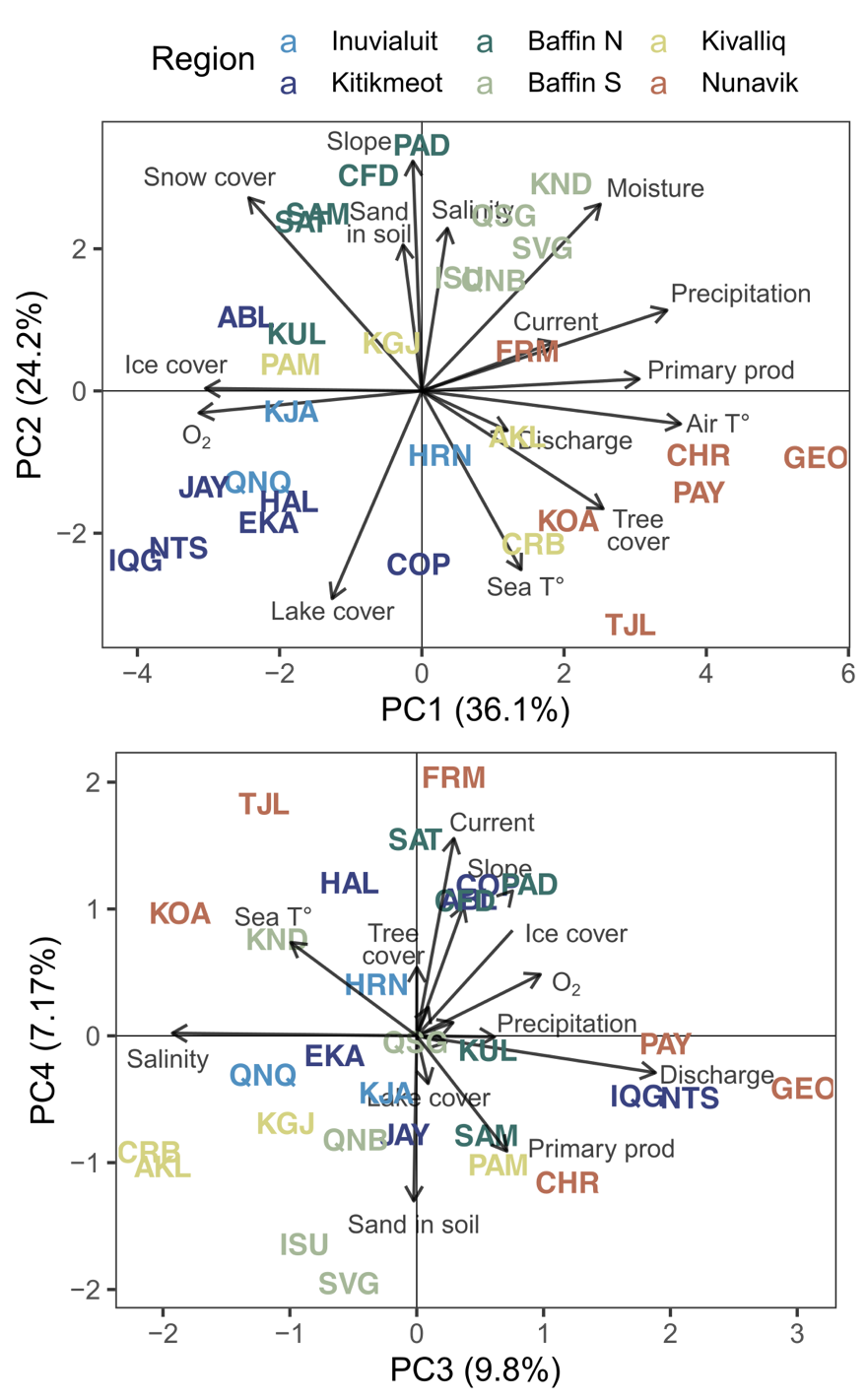

Fig S2: Principal component analysis for genotype likelihoods at 277,570 independent SNPs. a) Proportion (%) of variance explained by the 20 first axes. Individual loadings for b) PC3-4, c) PC 5-6, d) PC 7-8, and e) PC 9-10. 95% confidence interval ellipses are drawn around each sampling site and the percentage of variance explained by each axis is noted in parentheses.

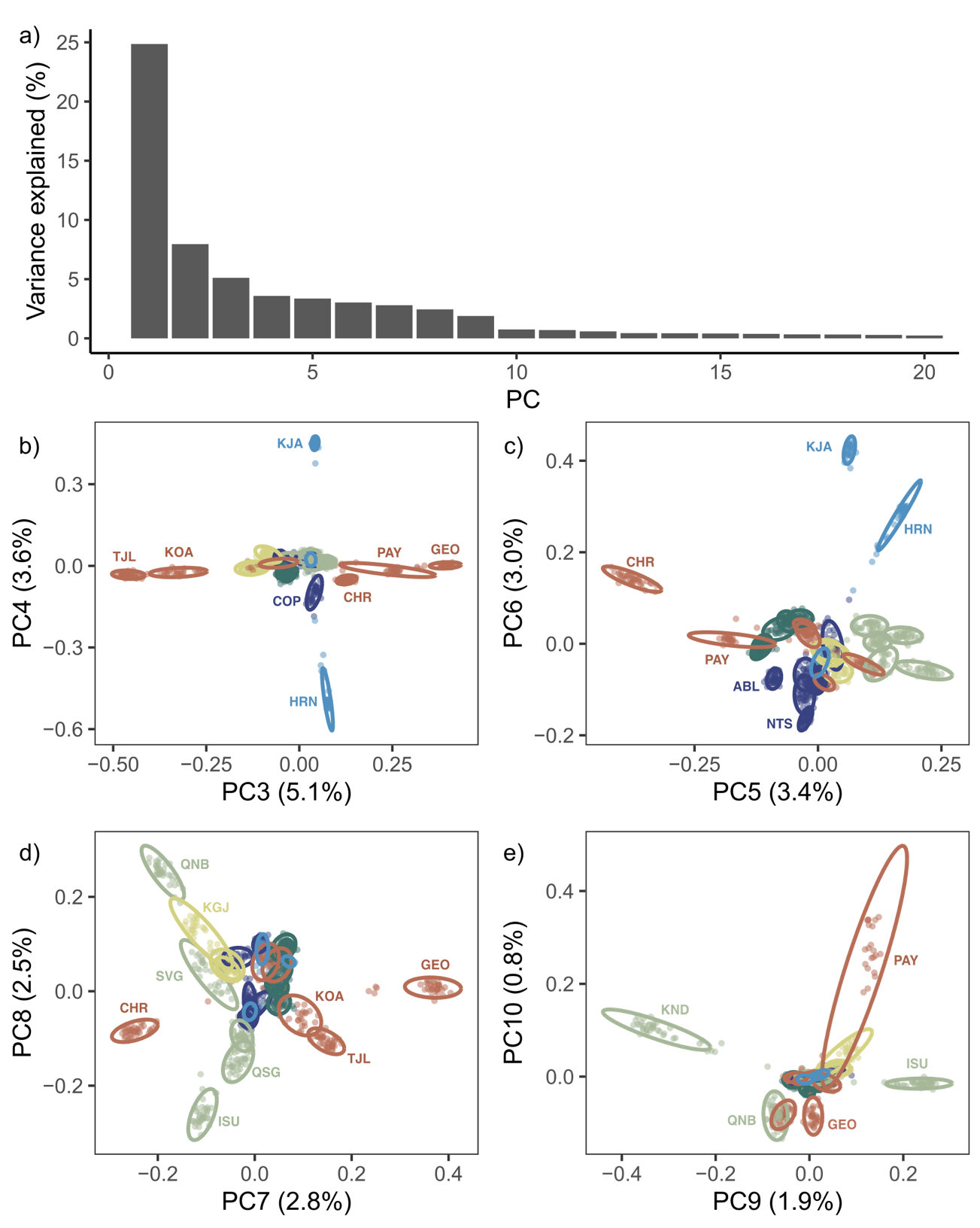

Fig S3 : a) Average best likelihood for 50 runs of NGSadmix with K ranging from 1 to 30, with standard deviation in error bars. b) ΔK estimated from the second-order rate of change in average likelihood divided by the standard deviation. c) Proportion of 50 runs in the major (black) and first minor (red) mode, as grouped by CLUMPAK with a 0.85 similarity threshold.

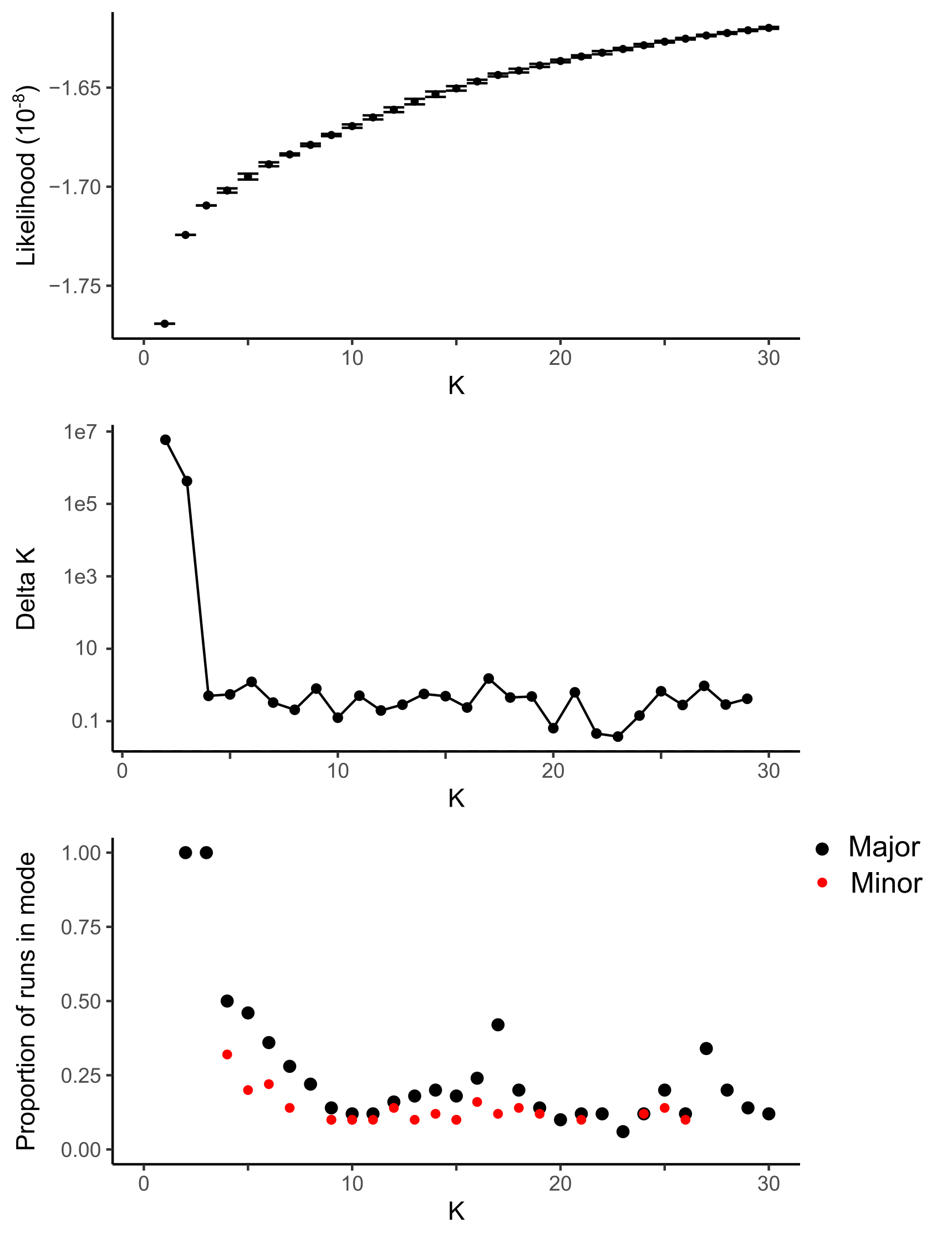

Fig S4 : Proportion of ancestry by individuals for K = 2 to 30 genetics clusters, CLUMPAK-averaged over replicate runs in the major mode (out of 50 runs). Individuals in all panels are ordered and aligned following the dominant genetic clusters in their sampling site for K = 7.

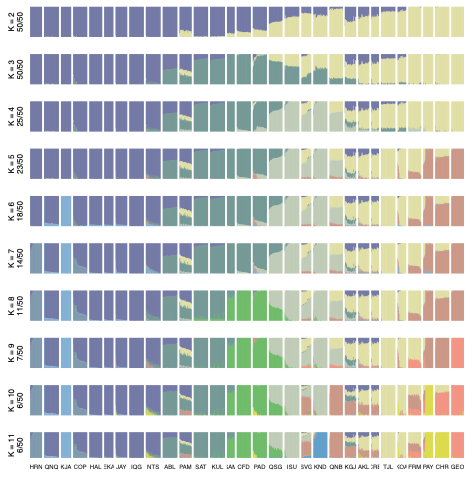

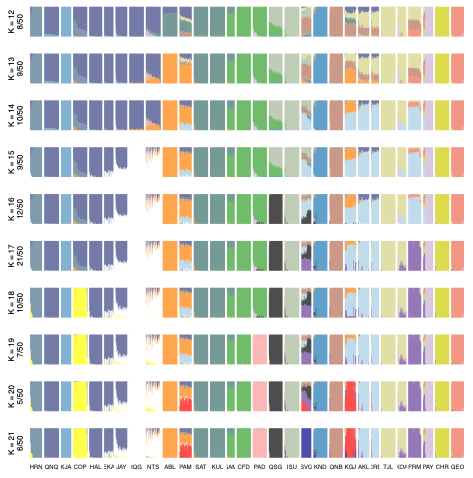

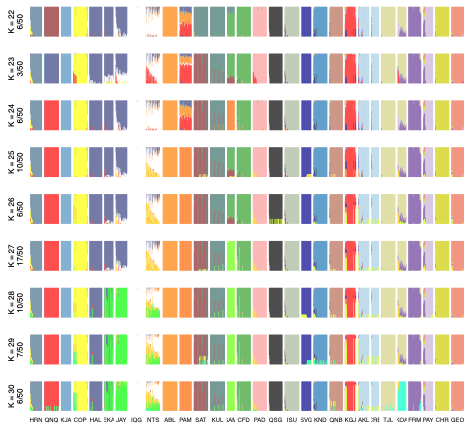

Fig S5 : Relation between residual Allele Frequency Difference (i.e. after correcting for ancestry distance) and difference in every environmental variables for a) all, b) northern, and c) southern population pairs. Linear regressions were fitted on scatter plots, and the r statistic from a partial Mantel test with marine distance as a covariable (as represented in color from blue to yellow) are displayed. Partial Mantel tests with p-values under 0.01 after 999 permutations are in red.

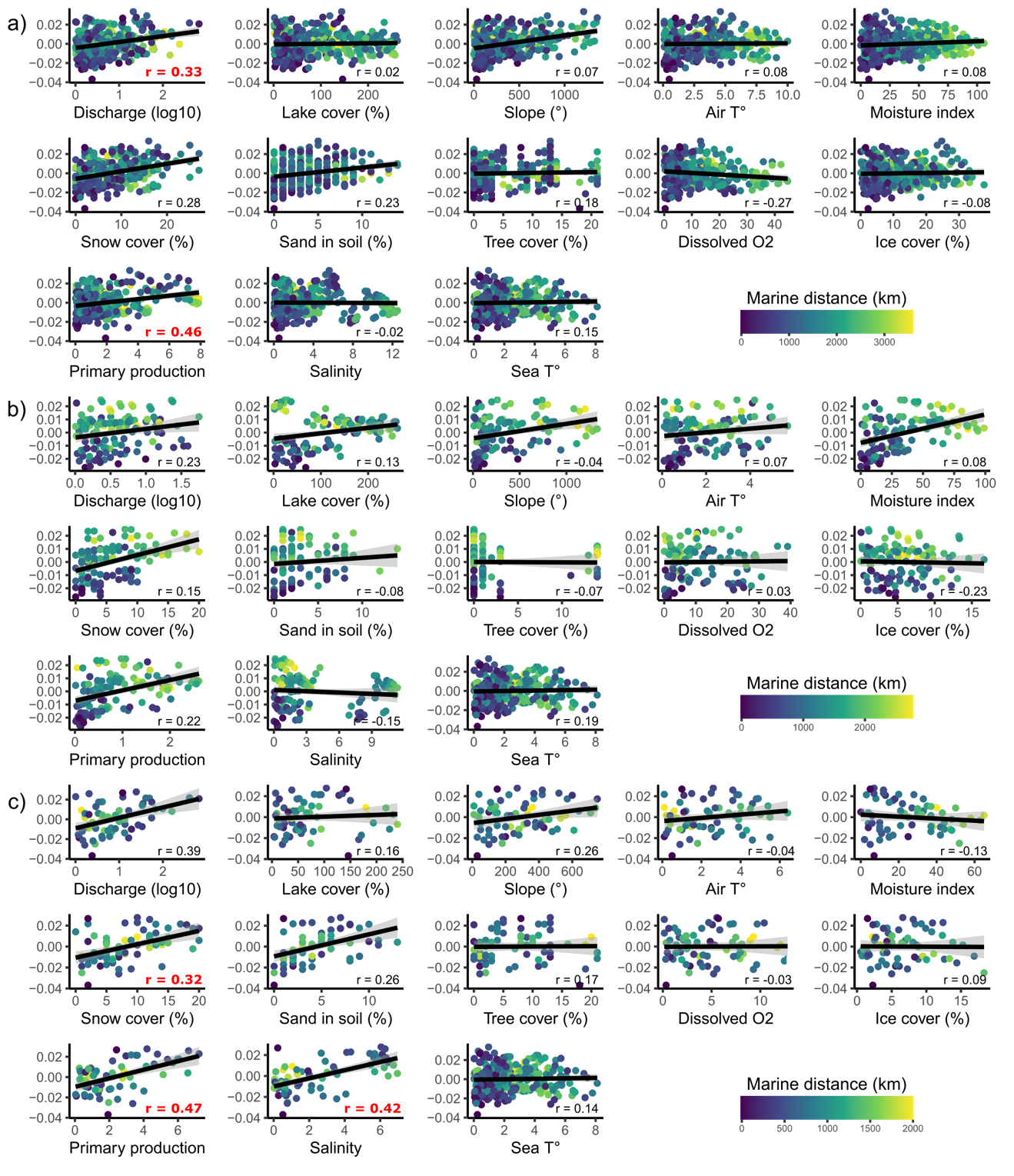

Fig. S6: Edge-specific effective migration rates (*w*) estimated from sampling allele frequencies in FEEMS with a smoothing parameter *λ* = a) 6.8, b) 21.5, and c) 146.8. Sampling sites were repositioned to the nearest node on a triangular grid (cell spacing ≈ 55 km). d) Average cross-validation error for *λ* ranging from 0.1 to 1000. *λ* values from local minima used in panel a-c are noted by letters instead of points.

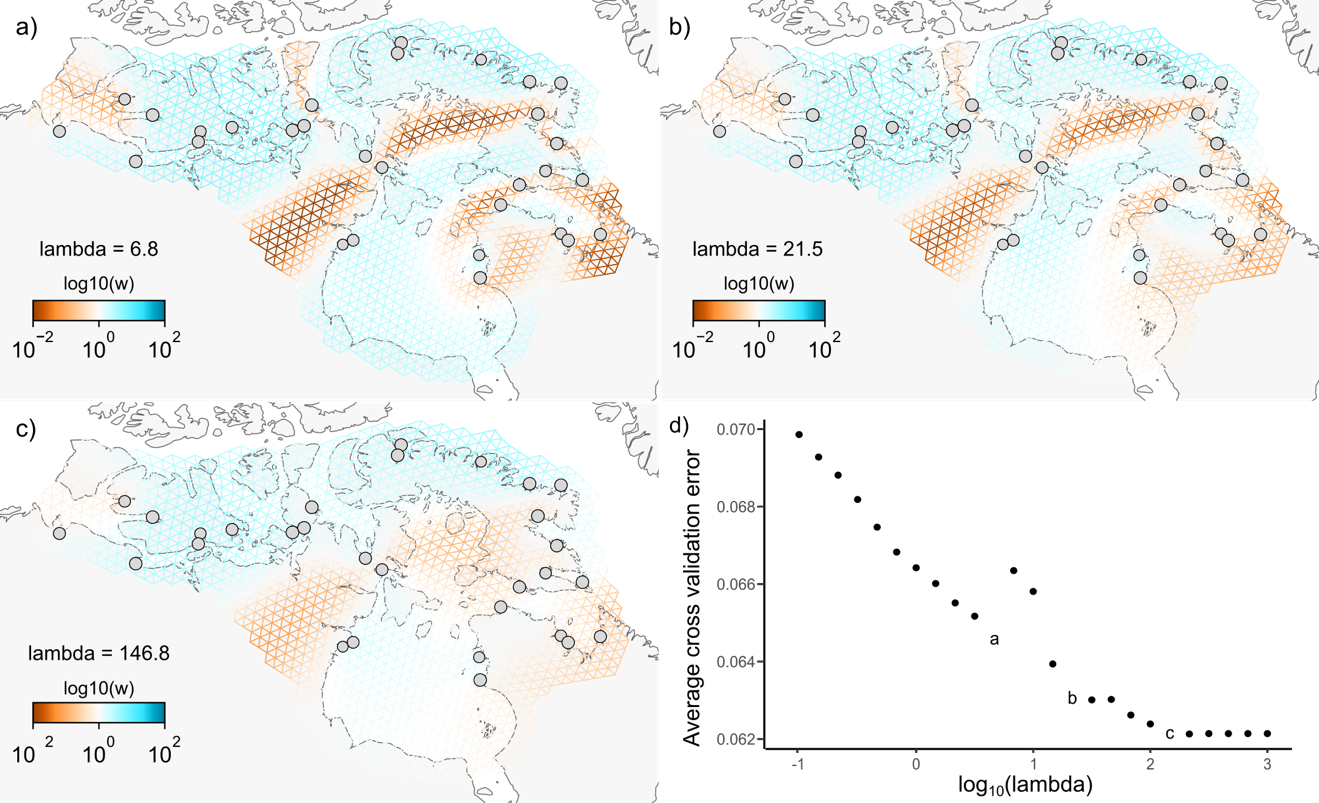

Figure S7: a) Support for selection scans or gene-environment associations on genes with at least 10 SNPs on 4 arbitrarily chosen linkage groups, in 5 panels regrouping methods: 1) XtX (Baypass) or pcadapt; 2) redundancy analysis axes (RDA); 3) partial RDA axes; and Bayes Factors (BF, Baypass) for 4) freshwater and 5) marine variables. Support at the gene level (empirical p-values) was derived from combined per-SNP summary statistics in a windowed Z analysis (WZA). Genes with the strong support (p < 0.001) for more than one type of test were highlighted with bigger diamonds than the rest (small dots). Shaded blue areas mark the postion of putative haplotype blocks identified in Dallaire et al. 2025. See Figure 5 for legend.

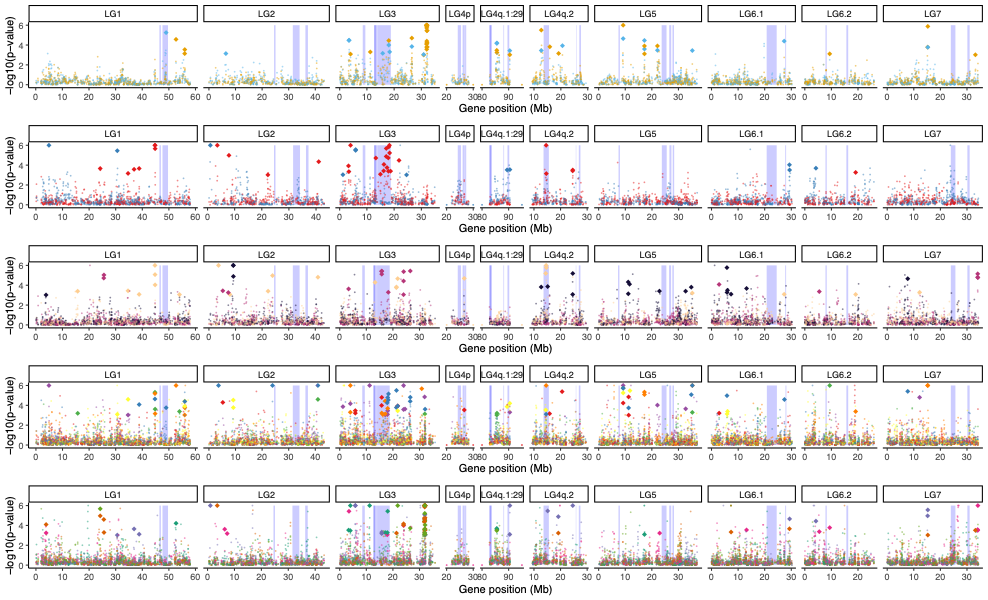

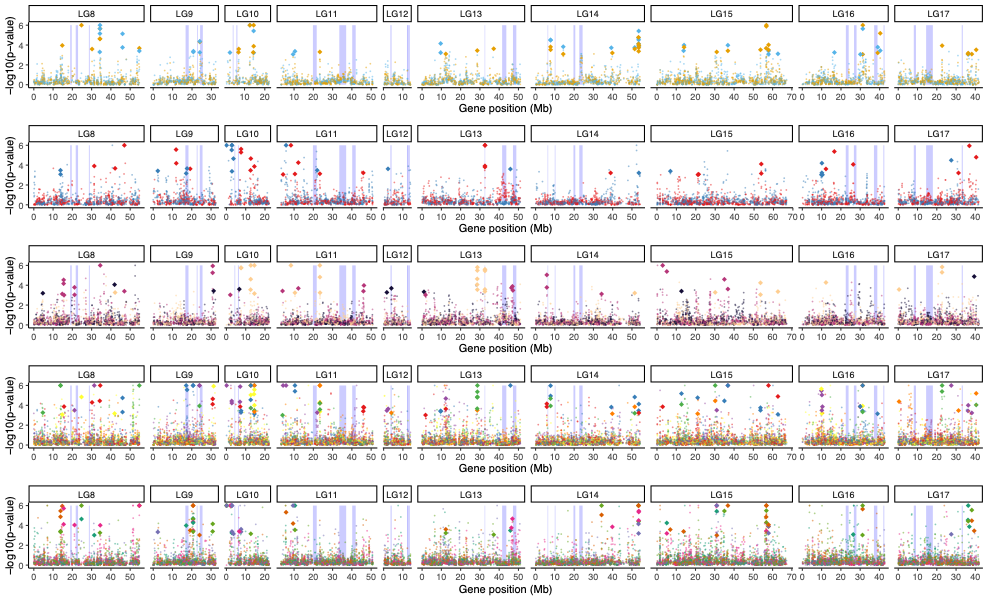

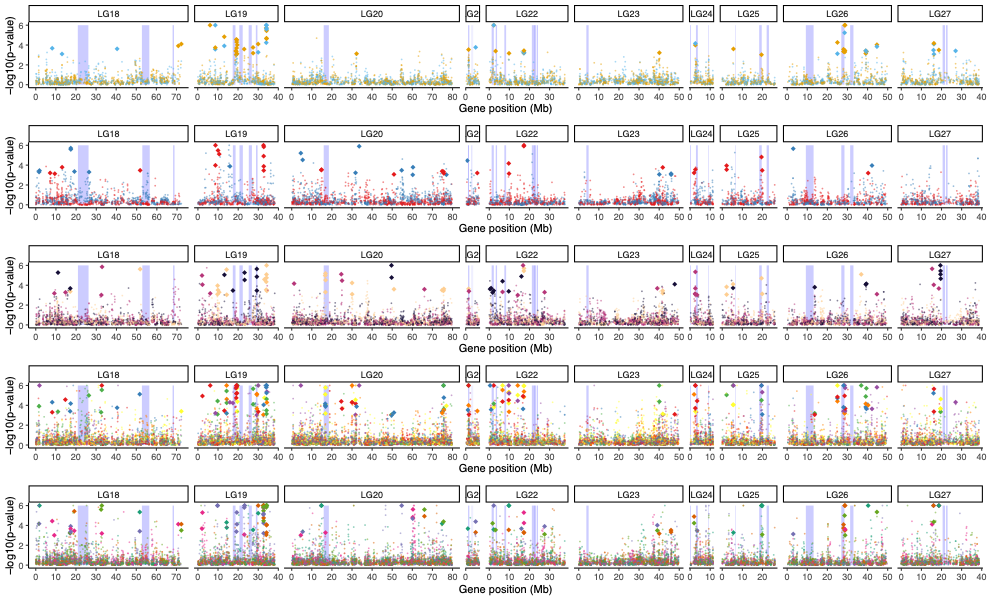

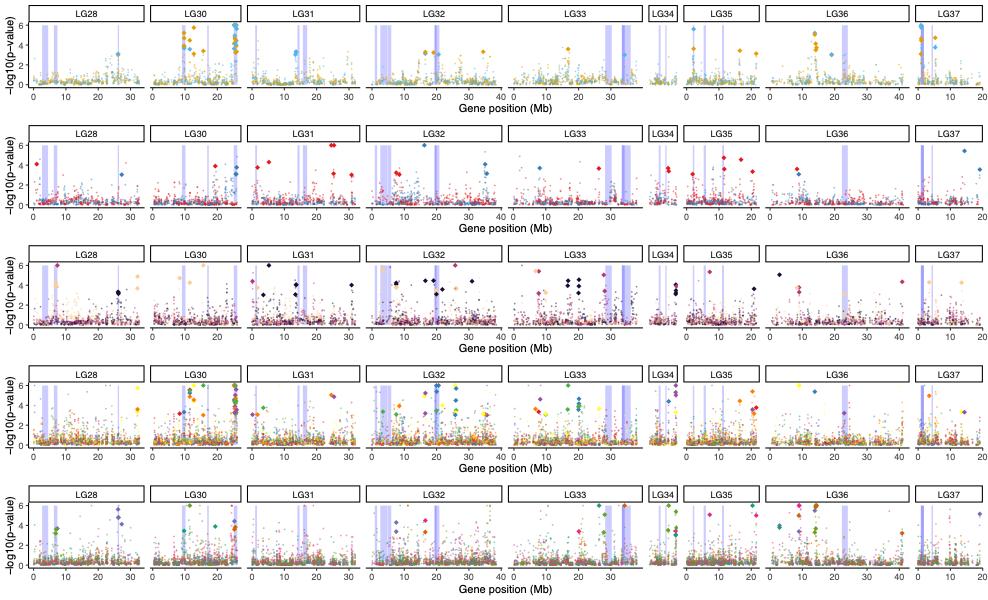
